# Oxidant-Resistant LRRC8A/C Anion Channels Support Superoxide Production by Nox1

**DOI:** 10.1101/2021.02.03.429614

**Authors:** Hyehun Choi, Jeffrey Rohrbough, Hong N. Nguyen, Anna Dikalova, Fred S. Lamb

**Affiliations:** Department of Pediatrics, Vanderbilt University Medical Center, Nashville, TN 37232

**Keywords:** Volume Regulated Anion Channel, Leucine Rich Repeat Containing 8, Tumor Necrosis Factor-α, Superoxide, NADPH Oxidase 1, Chloramine T, Oxidation

## Abstract

Tumor necrosis factor-α (TNFα) activates NADPH Oxidase 1 (Nox1) in vascular smooth muscle cells (VSMCs), producing superoxide (O_2_^•-^) required for subsequent signaling. LRRC8 family proteins A-E comprise volume-regulated anion channels (VRACs). The required subunit LRRC8A physically associates with Nox1, and VRAC activity is required for Nox activity and the inflammatory response to TNFα. LRRC8 channel currents are modulated by oxidants, suggesting that oxidant sensitivity and proximity to Nox1 may play a physiologically relevant role. In VSMCs, LRRC8C knockdown (siRNA) recapitulated the effects of siLRRC8A, inhibiting TNFα-induced extracellular and endosomal O_2_^•-^ production, receptor endocytosis, NF-κB activation, and proliferation. In contrast, siLRRC8D potentiated NF-κB activation. Nox1 co-immunoprecipitated with 8C and 8D, and co-localized with 8D at the plasma membrane and in vesicles. We compared VRAC currents mediated by homomeric and heteromeric 8C and 8D channels expressed in HEK293 cells. The oxidant chloramine T (ChlorT, 1 mM) weakly inhibited LRRC8C, but potently inhibited 8D currents. ChlorT exposure also greatly reduced subsequent current block by DCPIB, implicating external sites of oxidation. Substitution of the extracellular loop domains (EL1, EL2) of 8D onto 8C conferred significantly stronger ChlorT-dependent inhibition. 8A/C channel activity is thus more effectively maintained in the oxidized microenvironment expected to result from Nox1 activation at the plasma membrane. Increased ratios of 8D:8C expression may potentially depress inflammatory responses to TNFα. LRRC8A/C channel downregulation represents a novel strategy to reduce TNFα-induced inflammation.

**Key Points:**

- LRRC8A-containing anion channels associate with Nox1 and regulate superoxide production and TNFα signaling. Here we show that .LRRC8C and 8D also co-immunoprecipitate with Nox1 in vascular smooth muscle cells.
- LRRC8C knockdown inhibited TNFα-induced O_2_^•-^ production, receptor endocytosis, NF-κB activation, and proliferation while LRRC8D knockdown enhanced NF-κB activation. Significant changes in LRRC8 isoform expression in human atherosclerosis and psoriasis suggest compensation for increased inflammation.
- The oxidant chloramine-T (ChlorT, 1 mM) weakly (∼25%) inhibited 8C currents but potently (∼80%) inhibited 8D currents. Substitution of the two extracellular loop (EL) domains of 8D onto 8C conferred significantly stronger ChlorT-dependent inhibition.
- ChlorT also impaired current block by DCPIB, which occurs through interaction with EL1, further implicating external sites of oxidation.
- 8A/C channels most effectively maintain activity in an oxidized microenvironment, as is expected to result from Nox1 activity at the plasma membrane.

## INTRODUCTION

TNFα causes the production of extracellular superoxide (O_2_^•-^) by activating NADPH Oxidase 1 (Nox1) at the plasma membrane (Choi *et al*., 2016). O_2_^•-^ production continues within “signaling endosomes” formed as TNFα receptors are internalized (Miller *et al*., 2007). This reactive oxygen species (ROS) production is critical for inflammatory signaling in vascular smooth muscle cells (VSMCs) as demonstrated by the fact that interference with Nox1 activity or scavenging of ROS blocks TNFα signaling (Miller *et al*., 2007; Gimenez *et al*., 2016). NADPH oxidases transfer two electrons from cytoplasmic NADPH to molecular oxygen on the opposite side of an insulating membrane. Sustained Nox activity therefore requires ongoing charge compensation (DeCoursey *et al*., 2003). HV1 proton channels serve this role by operating in parallel to Nox2 in plasmalemmal and phagosomal membranes of white blood cells (DeCoursey, 2016). It has been speculated that another channel(s) provides charge compensation in other cell types for Nox1, but this has not been directly established (Miller *et al*., 2007; Lamb *et al*., 2009; Choi *et al*., 2016).

We recently demonstrated that <u>L</u>eucine <u>R</u>ich <u>R</u>epeat <u>C</u>ontaining 8A (LRRC8A), an essential component of Volume-Regulated Anion Channels (VRACs), is part of the multi-protein Nox1 signaling complex in VSMCs. LRRC8A physically associates with Nox1, and VRAC activity is required for full Nox1 activation in response to TNFα. Furthermore, targeted LRRC8A knockdown disrupts multiple aspects of the downstream inflammatory response to TNFα. (Choi *et al*., 2016). These findings are consistent with prior observations that VRAC inhibitors and siLRRC8A modulate VSMC proliferation (Qian *et al*., 2009; Liang *et al*., 2014; Lu *et al*., 2019), and suggest that LRRC8 channels play a critical role in TNFα signaling. The nature of the functional dependence of Nox1 on VRAC remains unclear, but the voltage-dependence and rectification properties of VRACs are appropriate to support a role in charge compensation.

VRACs are hexameric proteins (Deneka *et al*., 2018). LRRC8A must combine with at least one additional LRRC8 subunit (B,C,D,E) (Qiu *et al*., 2014; Voss *et al*., 2014; Pedersen *et al*., 2015) to produce active plasma membrane channels. Different subunit combinations yield currents with distinct biophysical properties in terms of single channel conductance, open probability and ion selectivity (Voss *et al*., 2014; Syeda *et al*., 2016). Heterologous co-expression of LRRC8A/C, A/D, and A/E isoforms in *Xenopus* oocytes resulted in currents with differential sensitivity to the oxidant chloramine T, which activated A/E channel currents, but inhibited A/C and A/D channel currents (Gradogna *et al*., 2017). The physical association of Nox1 with redox-sensitive channels capable of regulating Nox1 activity and ROS production raises the possibility of reciprocal functional interdependence between these two proteins. Redox-dependent modification of LRRC8 VRACs could provide highly localized feedback regulation of Nox1 activity.

We sought to determine which LRRC8 channel subtypes associate with Nox1 and how they support oxidase activity and TNFα signaling in VSMCs. siRNA-mediated knockdown of LRRC8C had an anti-inflammatory impact on TNFα signaling similar to that previously identified for siLRRC8A (Choi *et al*., 2016), while siLRRC8D knockdown had significant but opposing effects. Neither siLRRC8B nor siLRRC8E significantly altered TNFα signaling. siLRRC8C also reduced TNFα-induced extracellular O_2_^•-^ production, demonstrating an impact on Nox1 activity, while siLRRC8D did not. VRAC currents derived from LRRC8C expression were relatively resistant to ChlorT-induced inhibition compared to LRRC8D channel currents, which were potently inhibited. Thus, LRRC8A/C VRACs appear to be selectively suited to support sustained Nox1 activity and pro-inflammatory signaling. LRRC8 isoform expression and differential regulation by oxidation may represent important mechanisms for modulation of the inflammatory response to TNFα, as well as other pro-inflammatory signaling molecules that activate Nox1.

## MATERIALS AND METHODS

### Reagents

All chemicals and enzymes unless otherwise noted were obtained from Sigma-Aldrich (St. Louis, MO).

### Cell culture

Primary murine aortic VSMC were isolated from C57/BL6 mice as described previously (Choi *et al*., 2015). The cells were maintained in Dulbecco’s Modified Eagle’s Medium (DMEM; Life Technologies, Grand Island, NY) supplemented with 10% fetal bovine serum (FBS), 1% penicillin/streptomycin, 1X minimum essential medium non-essential amino acids, 1X vitamin, and 20 mM HEPES. Human aortic VSMCs were purchased from Cell Applications (San Diego, CA) and maintained in human SMC growth medium (#311-500, Cell Applications).

HEK293T cells were obtained from the American Tissue Culture Collection. HEK293 cells lacking LRRC8A were the generous gift of Dr. Rajan Sah (Washington University, St. Louis MO). These cells were maintained at 37° C in 5% CO_2_ in Dulbecco’s modified Eagle’s medium (DMEM), supplemented with 10% fetal bovine serum (FBS, Hyclone) and 0.1% Pen/Strep (25U/ml, Life Technologies).

### Polymerase Chain Reaction

Qualitative comparisons of the expression of LRRC8 isoforms were made by reverse transcription-PCR analysis of mRNA isolated from VSMCs and HEK293 cells. The primers employed are provided in **Table 1**.

**Table 1.**
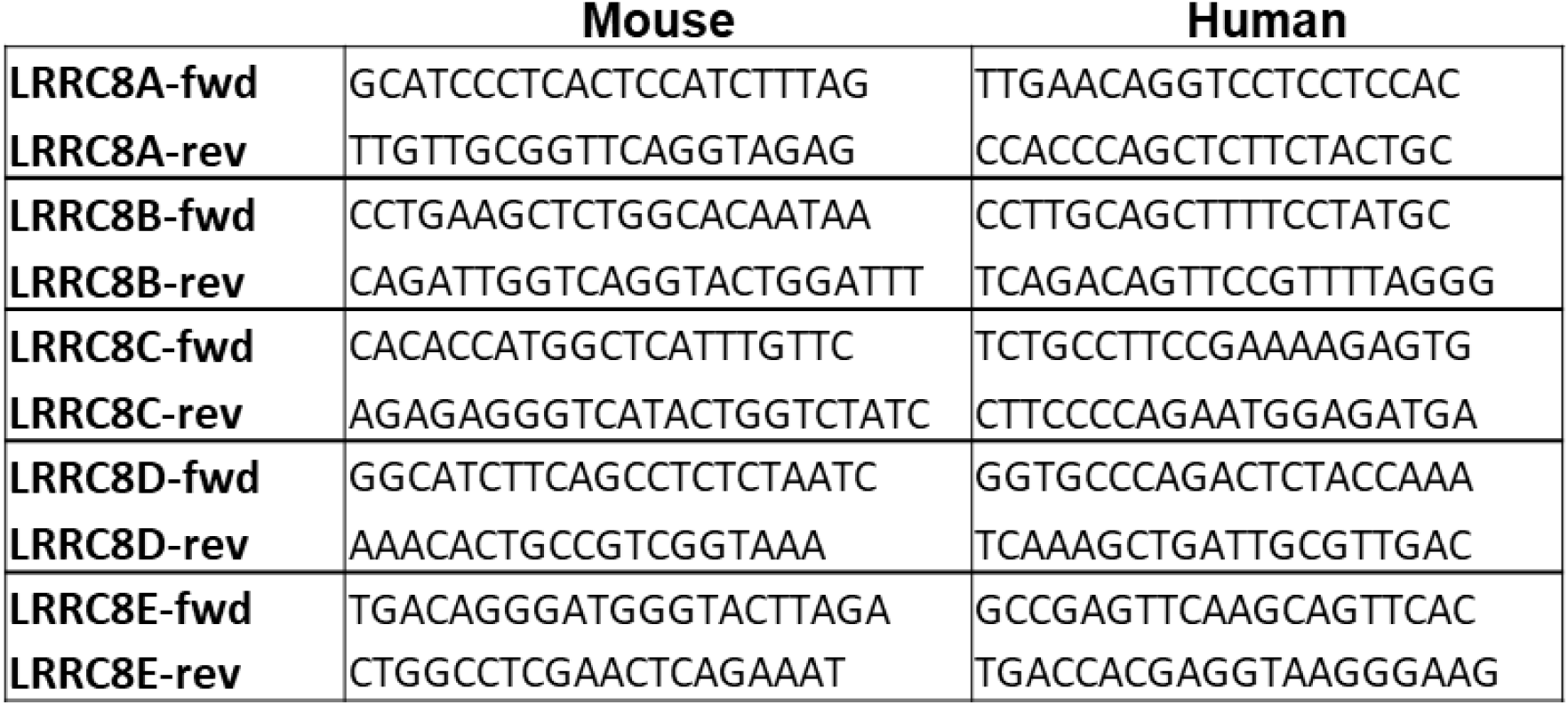

### Plasmid Modification and Expression

Wild-type LRRC8A (RC508632), C (RC203642), and D (RC222603) were obtained from Origene Technologies, and E constructs was obtained from ViGene Biosciences (CH871303) Chimeric constructs in which the intracellular loop of LRRC8A was inserted into LRRC8C, D or E were kindly provided by Dr. Kevin Strange (Novo Biosciences and Vanderbilt University). Modifications of these plasmids were made using the QuikChange Lightning site-directed mutagenesis kit (Agilent). The C-termini were tagged with mGFP or mCherry to facilitate protein localization. Substitution of the two extracellular loops were made as detailed in Results. Plasmids were transfected using Lipofectamine 2000 (DMEM + 1.5-2.5 µg/µl cDNA, 2.5% lipofectamine 2000, +10% FBS). For co-transfections of LRRC8A and a second isoform a cDNA ratio of 3:1 to 4:1 (nonA:A) was used.

### siRNA transfection

siRNA (negative control, LRRC8A, B, C, D, E) were purchased from Dharmacon (Lafayette, CO). siRNA (100 nM) was incubated with Lipofectamine 2000 (Life Technologies) in serum-free medium for 15 min. The resultant complex of siRNA-Lipofectamine 2000 was added to cells in DMEM containing 5% FBS and then maintained for 3 days before performing experiments.

### NF-κB activity

VSMC were infected following 2 days of siRNA transfection with replication-deficient adenovirus expressing a luciferase reporter driven by NF-κB transcriptional activation for 24 hours in DMEM containing 5% FBS followed by exposure to TNFα (10 ng/mL) in serum-free DMEM for 6 hours. Luciferase activity (relative light units) was quantified according to the manufacturer’s protocol (Promega, Madison, WI) and normalized to protein concentration (BCA protein assay).

### Immunoprecipitation (IP)

Cells were stimulated with TNFα (10 ng/mL, 10 min) then lysed (50mM Tris base, 150mM NaCl, 1mM EDTA, 10% glycerol, 1mM DTT, 1% Triton X-100, 0.1% Na-DOC, 0.1% SDS, 10mM β-glycerophosphate, 20mM para-nitrophenyl phosphate, 2mM sodium pyrophosphate, 1mM Na_3_VO_4_, 5mM NaF, 10 µg/ml aprotinin, and 1 mM phenylmethylsulfonyl fluoride (PMSF) at pH 7.4) for 1 hour with nutation at 4°C, and centrifuged for 30min at 20,000g. Supernatants were pre-cleared with protein-G sepharose beads for 1h at 4°C and cleared-supernatants were incubated with antibody (2 µg) for 1.5h, then incubated with protein-G sepharose for 1h. Beads were washed with lysis buffer, resuspended in SDS sample buffer, boiled and the associated proteins were then analyzed by western blot. Antibodies for IP were used as follows; Nox1 (sc-5821, Santa Cruz), LRRC8C (HPA029347, Sigma-Aldrich), LRRC8D (HPA014745, Sigma-Aldrich).

### Immunofluorescence

HEK293T and human aortic VSMCs were grown on coverslips and fixed in 4% paraformalde for 15min at room temperature. For HEK293T cells the coverslips were coated with 20 µg/ml polyethyleneimine (50-100,000 MW) to facilitate adherence. Cells were permeabilized with 0.5% Triton-X 100 and blocked with 1% BSAprior to incubation with primary and secondary antibody (1:100) (Nox1 (Santa Cruz, sc-5821), (LRRC8D (Santa Cruz, sc-515070)) at 37°C for 1h in the dark. They were then mounted in ProLong Gold (ThermoFisher). Images were obtained on a Leica SPE confocal microscope.

### Western blot analysis

Cells were stimulated with TNFα (10ng/mL). Protein extracts (40 μg) were separated by electrophoresis on a polyacrylamide gel (10%) and transferred to nitrocellulose membranes. Nonspecific binding was blocked with Odyssey blocking buffer (LI-COR Biosciences, Lincoln, NE) for 1 hour at room-temperature. Membranes were incubated with primary antibodies overnight at 4°C. Antibodies included: Tubulin (Vanderbilt Antibody Core), LRRC8A (#A304-175A, Bethyl Laboratories, Montgomery, TX), LRRC8C (HPA029347, Sigma-Aldrich), LRRC8D (HPA014745, Sigma-Aldrich), PCNA (05-347, Millipore). Signals were developed using fluorescent secondary antibodies with the Odyssey Imaging System (LI-COR Biosciences, Lincoln, NE) and quantified densitometrically.

### Detection of TNFα receptor endocytosis

Cells were grown on coverslips in six well plates and incubated with siRNA for 3 days. Human TNFα conjugated with biotin (R&D Systems) was incubated with FITC-labeled avidin (Life Technologies) for 1 h at 4°C, and then exposed to cells for 2 h at 4°C. After washing with cold media, cells were warmed to 37°C for 15 min. Cells were then fixed in 3.7% formaldehyde and nuclear counterstaining was performed with TO-PRO-3 Iodide (Life Technologies) for 5 min. Cover-slides were mounted with ProLong Gold anti-fade reagent (Life Technologies). Cells were imaged by Leica SPE confocal laser microscope and the intracellular punctate FITC signal quantified using ImageJ software.

### Superoxide detection

Extracellular and endosomal O_2_^•-^ was quantified using the membrane-impermeable electron spin resonance (ESR) probe 1-Hydroxy-2,2,6,6-tetramethylpiperidin-4-yl-trimethylammonium chloride (CAT1H; Enzo Life Sciences). Cells were incubated with Krebs/HEPES containing CAT1H (0.5 mmol/L) and TNFα (10 ng/mL) for 20 minutes then washed, scraped and snap frozen and placed in an ESR finger Dewar under liquid nitrogen. ESR spectra were recorded from the cell samples (endosomal) or from the Krebs buffer (extracellular) using the following settings: field sweep, 80 G; microwave frequency, 9.39 GHz; microwave power, 2 mW; modulation amplitude, 5 G; conversion time, 327.68 ms; time constant, 5242.88 ms; 512 points resolution; and receiver gain, 1×10^4^. The ESR signal was normalized to protein concentration.

### Gene Expression in Human Disease

We utilized the GTExPortal (gtexportal.org) to compare LRRC8 isoform mRNA expression among 54 different tissues in samples from hundreds of individuals. GEO2R was used to analyze microarray data from the NIH Gene Expression Omnibus (GEO datasets, www.ncbi.nlm.nih.gov) and assess LRRC8 isoform expression in human biopsy samples from control patients and individuals with inflammatory aortic disease (GSE57691) or psoriasis (GSE13355).

### Electrophysiology

HEK 293T cells were seeded into 6- or 12-well plates for > 2 hrs and transfected overnight with plasmid. Whole-cell patch clamp recordings were made at room temperature (21-22 °C) from freshly trypsinized and dissociated cells collected on the 1^st^ day (18-24 hrs) or 2^nd^ day (42-48 hrs) post transfection. LRRC8 protein expression level and distribution was documented in all recorded cells by visualizing mGFP- and/or mCherry fluorescence on an Olympus IX71 microscope (60X objective), equipped with a Hamamatsu C10600 digital camera and Till Photonics Oligochrome and image acquisition/analysis hardware and software (Hunt Optics & Imaging, Inc, Pittsburgh, PA). Isotonic external recording saline (330 mosm/L, pH 7.4) contained (in mM): 130 NaCl, 1.8 MgCl_2_, 1.8 CaCl_2_, 10 HEPES, ∼70 Mannitol, 1-2 NaOH. Pipette saline (315 mosm/L, pH 7.2) contained (in mM): 120mM CsCl, 4 TEACl, 2 MgCl_2_, 5 Na_2_ATP, 10 HEPES, ∼35 Mannitol, 1-2 CsOH, 1.186 CaCl_2_, 5 EGTA (estimated free Ca^2+^ of 61 mM using WEBMAXC). Final solution osmolality was adjusted by addition of 1M mannitol, measured with a Precision Systems mOsmette osmometer (Natick, MA). Mannitol was excluded from the hypotonic external saline (250-255 mosm/L). DCPIB (30 μM; Tocris Bioscience, Bristol UK) was added daily from freshly thawed 30 mM (DMSO) stock aliquots. Chloramine-T (chlorT, 0.25-1.0 mM) and TCEP were added from 0.5M stock solutions. DTT (10 mM) was directly added to salines on the day of use. The recording chamber volume was maintained at ∼500 µl and superfused constantly at ∼1.2 ml/min. Solution changes were effected using an Automate ValveBankII system (Berkeley, CA) with a multiport manifold that delivered new solution to the recording chamber with a latency of 8-10s.

Amplified whole-cell currents were low-pass filtered at 2 kHz and sampled at 10 kHz using a Molecular Devices Axopatch 200B amplifier driven with a pClamp 10 interface (Molecular Devices, Sunnyvale, CA). Pipette resistances were typically 2.0-3.0 MΩ. Fast pipet capacitance was compensated following gigaseal formation. Cell capacitance (C_m_; 20 + 1 pF) and series resistance (R_S_; 6.6 + 0.2 MΩ) were measured using the Clampex 10 membrane test utility while applying a +20 mV test pulse from a holding potential (V_H_) of -40 mV. 65%-70% R_S_ compensation (75% prediction) was applied. Current-voltage (I-V) relationships were recorded for 100 ms test pulses (−100 to +120 mV, in 20 mV increments) applied from a prepulse level of 0 mV. Current amplitude was measured 2 ms after initiation of the voltage pulse. Oxidant effects were assessed while recording current responses to 250 ms voltage ramps (−100 to +120 mV) at 30s intervals. Ramp current values at +120 mV applied potential were quantified and reported. Current-voltage (I-V) plots are corrected for a 5 mV liquid junction potential measured between the pipette and isotonic bath saline. Current density is expressed as pA/pF. Zero-current level is indicated in displayed traces by a dashed line.

### Statistical analysis

Statistical calculations and comparisons were made with Excel (2013) and Graphpad Prism 5 analysis software (GraphPad Software, San Diego, CA). Values are reported as mean ± standard error of the mean (SEM). N represents the number of independently performed experiments in cultured cells, or the number of recorded cells in electrophysiological experiments. Plots were generated using Graph Pad Prism. For quantification of biochemical results, statistical differences were assessed by unpaired *t*-test, one sample *t*-test (control hypothetical value is 1) or one-way ANOVA. *Post hoc* comparisons were performed using Newman-Keuls analysis to compare all groups. Statistical significance reported for GEO gene expression data represent corrected P values for between group differences as calculated by the GEO2R software based on log_2_ transformed data. Comparison of ratios of LRRC8C to 8D expression in psoriasis were made after 2^X^ transformation of the data followed by one-way ANOVA and a non-parametric Kruskal-Wallis test. Ratio T-tests were applied to transformed VRAC current amplitude data to assess the significance of changes in paired measurements in control vs. oxidant conditions. The Mann-Whitney nonparametric test was used for other two-way comparisons. A *P* value less than 0.05 was considered to be statistically significant.

## RESULTS

### LRRC8C isoform supports TNFα –dependent inflammatory response in VSMCs

Messenger RNA was detected by PCR in murine VSMCs for all five members of the LRRC8 family (isoforms A through E). LRRC8C and 8D mRNA were the most abundant, while 8E mRNA was the least abundant (Fig. 1A). Efficacy of siRNA was verified by RT-PCR estimation of mRNA levels, which confirmed a pronounced knockdown of each isoform (Fig. 1B). To explore isoform-dependent roles in inflammatory signaling, we first assessed TNFα-mediated NF-κB activation. Consistent with our previous work (Choi *et al*., 2016), siLRRC8A reduced NF-κB activation by 49 ± 3%. NF-κB activation was also decreased by LRRC8C siRNA (31 ± 5%; N.S. vs. siLRRC8A) but was significantly potentiated by siLRRC8D (47 ± 29% increase; Fig. 1C). Knockdown of LRRC8B and LRRC8E had no significant effect on TNFα-mediated NF-κB activation. Consequently, we focused our subsequent work on the C and D isoforms. The nuclear protein PCNA (<u>P</u>roliferating <u>C</u>ell <u>N</u>uclear <u>A</u>ntigen) is involved in DNA replication and is an effective marker of proliferation. TNFα caused an increase in PCNA expression in murine VSMCs after 24 hrs in the presence of control siRNA (21 ± 5.7 % above control; Fig. 1D, E). Knockdown of LRRC8C insignificantly reduced PCNA abundance under control conditions (P = 0.084, n = 4), but siLRRC8C prevented the TNFα–induced increase in PCNA (P < 0.05 vs. siControl + TNFα; Fig. 1E). In contrast, siLRRC8D had no significant effect on either resting or stimulated PCNA levels.

**Figure 1.**
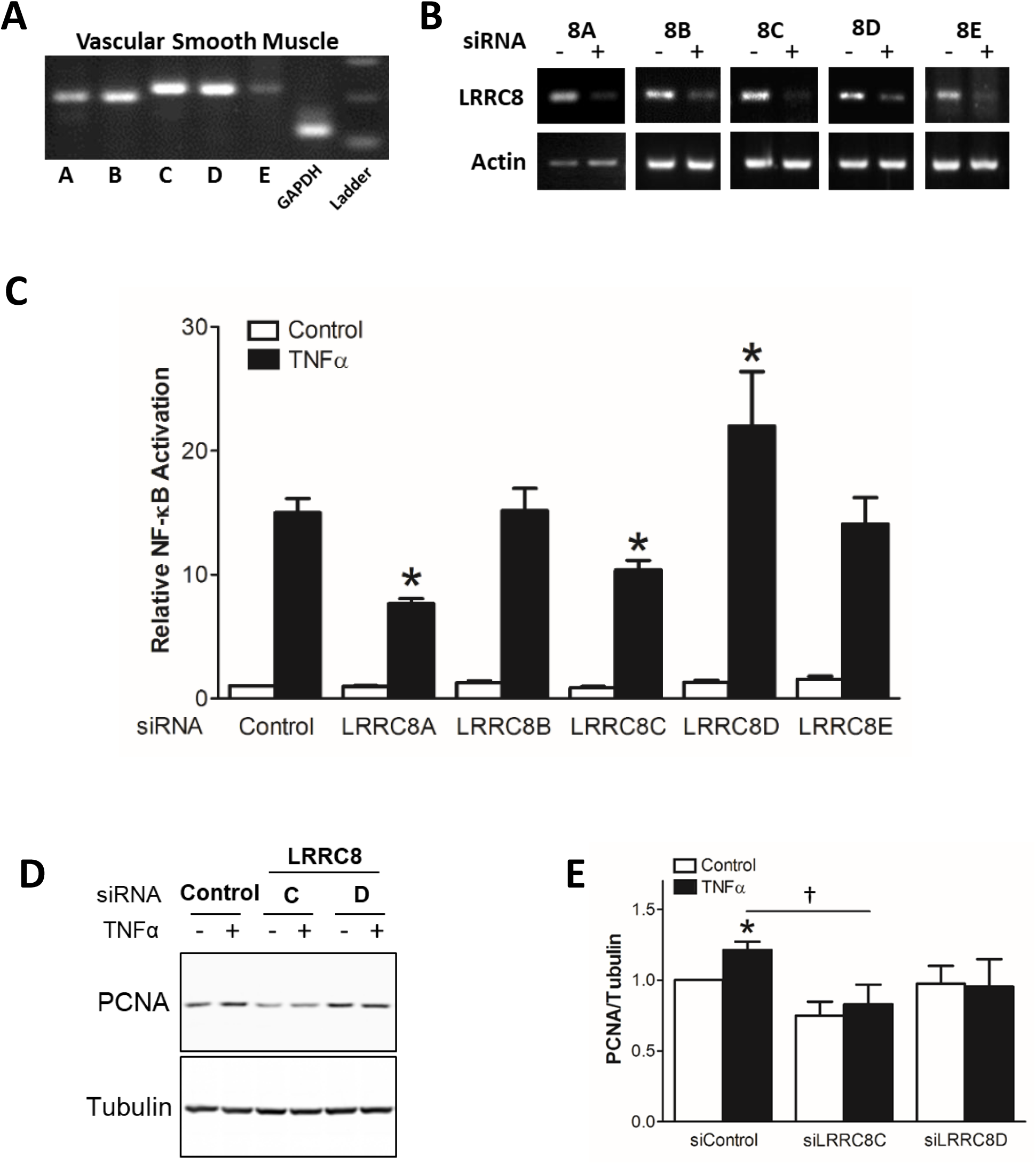
***A***, All LRRC8 isoforms (LRRC8A-E) are expressed in cultured murine aortic VSMCs. RT-PCR was performed with cDNA derived from mRNA obtained from cultured VSMCs. ***B***, siRNA knockdown of mRNA for LRRC8 isoforms was detected by RT-PCR. ***C***, NF-κB activation 6 hrs after exposure to TNFα in siRNA treated cells. Data are expressed as fold change compared to Control siRNA exposed cells that were not treated with TNFα (*\* p* < 0.05 compared to siControl cells exposed to TNFα; n = 10). ***D and E***, Activation of VSMC proliferation by TNFα is reflected by increased PCNA expression. This effect was lost in cells with reduced expression of LRRC8C or LRRC8D (\**P* < 0.05 compared to siControl cells not exposed to TNFα; † *P* < 0.05; n = 4).

Our previous work demonstrated that TNFα-induced NF-κB activation in VSMC is Nox1-dependent, and that LRRC8A physically associates with Nox1 by immunoprecipitation and immunohistochemistry results (Choi *et al*., 2016). Immunoprecipitation of Nox1 and western blotting for LRRC8C and 8D revealed associations between Nox1 and both proteins that were not significantly altered by treatment of the cells with TNFα (Fig. 2A). Immunostaining confirmed that LRRC8D co-localized with Nox1 at both the plasma membrane and in intracellular vesicles (Fig. 2B). Human aortic VSMCs were used for these experiments as they provided superior discrimination of these proteins by immunohistochemistry compared to murine VSMCs. Immunostaining was validated by comparing localization of a fluorescent tag on heterologously expressed proteins with the signal generated by immunostaining of that protein in HEK293 cells (Fig. 2C). Since the anti-LRRC8C and anti-Nox1 antibodies that demonstrated effective immunostaining were both raised in goat, we were unable to compare the localization of LRRC8C to Nox1.

**Figure 2.**
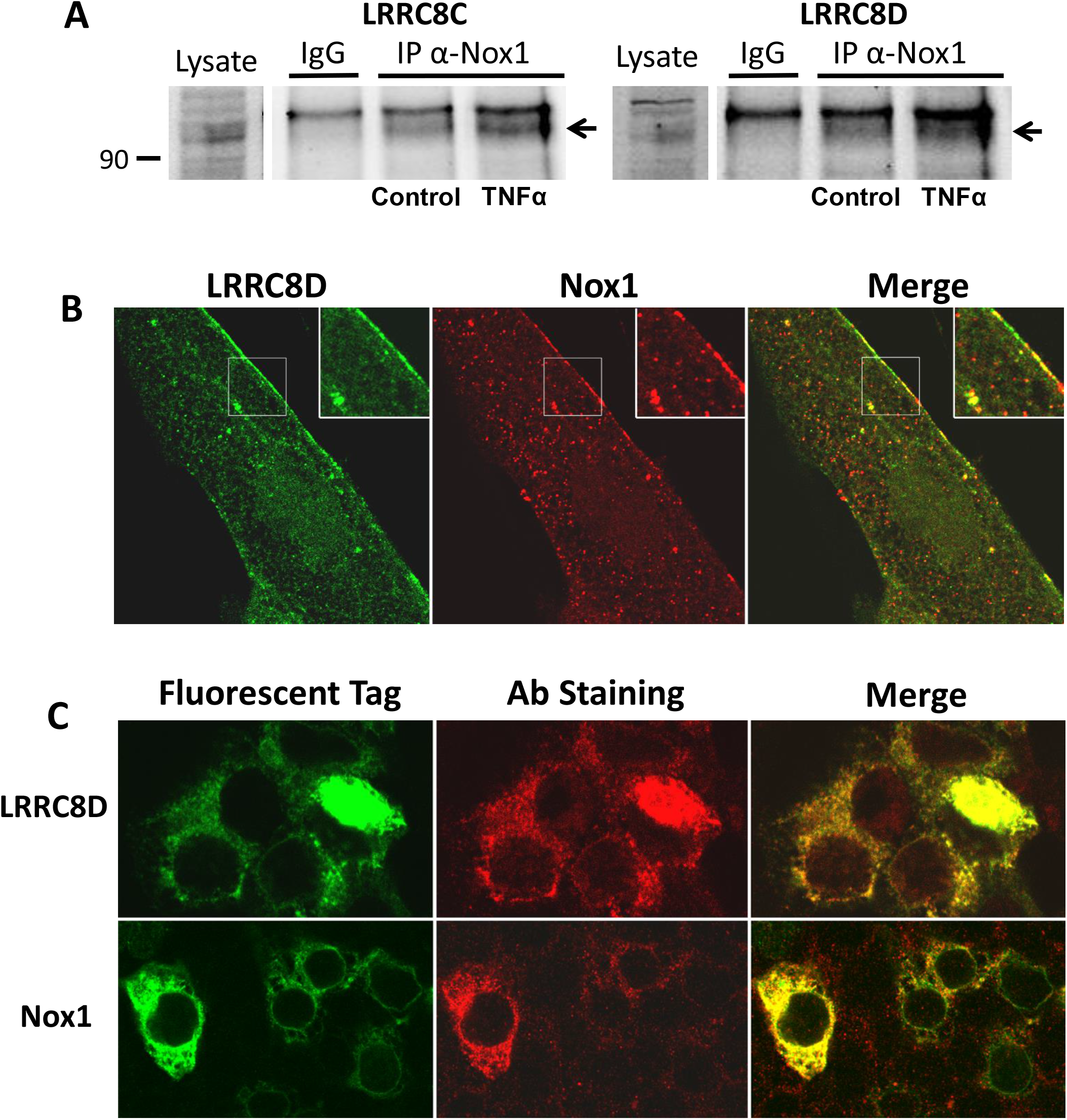
***A***, Control IgG or anti-Nox1 Ab were used to immunoprecipitate VSMC LRRC8C (left) or LRRC8D (right) proteins under control conditions or following a 10 min exposure to TNFα. Western blotting was subsequently performed on whole cell lysates (Lysate) or the immunoprecipitated proteins. Anti-Nox1 pulled down both LRRC8 proteins but this association was not enhanced by exposure to TNFα (n = 3). ***B***, Immunostaining of cultured human VSMCs with anti-LRRC8D and anti-Nox1 reveals co-localization both at the plasma membrane and in intracellular vesicles (insets shown at 2X magnification). ***C***, HEK293 cells heterologously expressing GFP-tagged LRRC8D (top) or Nox1 (bottom) immunostained with anti-LRRC8D or anti-Nox1 (middle panels in red), to validate antibody specificity. GFP localization and antibody labeling are strongly colocalized (merged panels, right).

TNFα-induced O_2_^•-^ production by Nox1 occurs at both the plasma membrane and within endosomes (Miller *et al*., 2007). Extracellular O_2_^•-^ production is primarily dependent on Nox1, as reflected by the markedly reduced O_2_^•-^ signal detected by the membrane-impermeable CAT1H spin trap in Nox1 null cells (Choi *et al*., 2019). We previously showed that siLRRC8A significantly reduced the quantity of extracellular O_2_^•-^ produced by TNFα-stimulated VSMCs (Choi *et al*., 2016). Here we quantified both extracellular and endosomal O_2_^•-^ production using CAT1H in control and siRNA-treated murine VSMCs cells. The abundance of O_2_^•-^ increased in both compartments following exposure to TNFα. Knockdown of LRRC8C significantly reduced TNFα-induced O_2_^•-^ production at both sites, while siLRRC8D had no effect (Fig. 3A, B).

**Figure 3.**
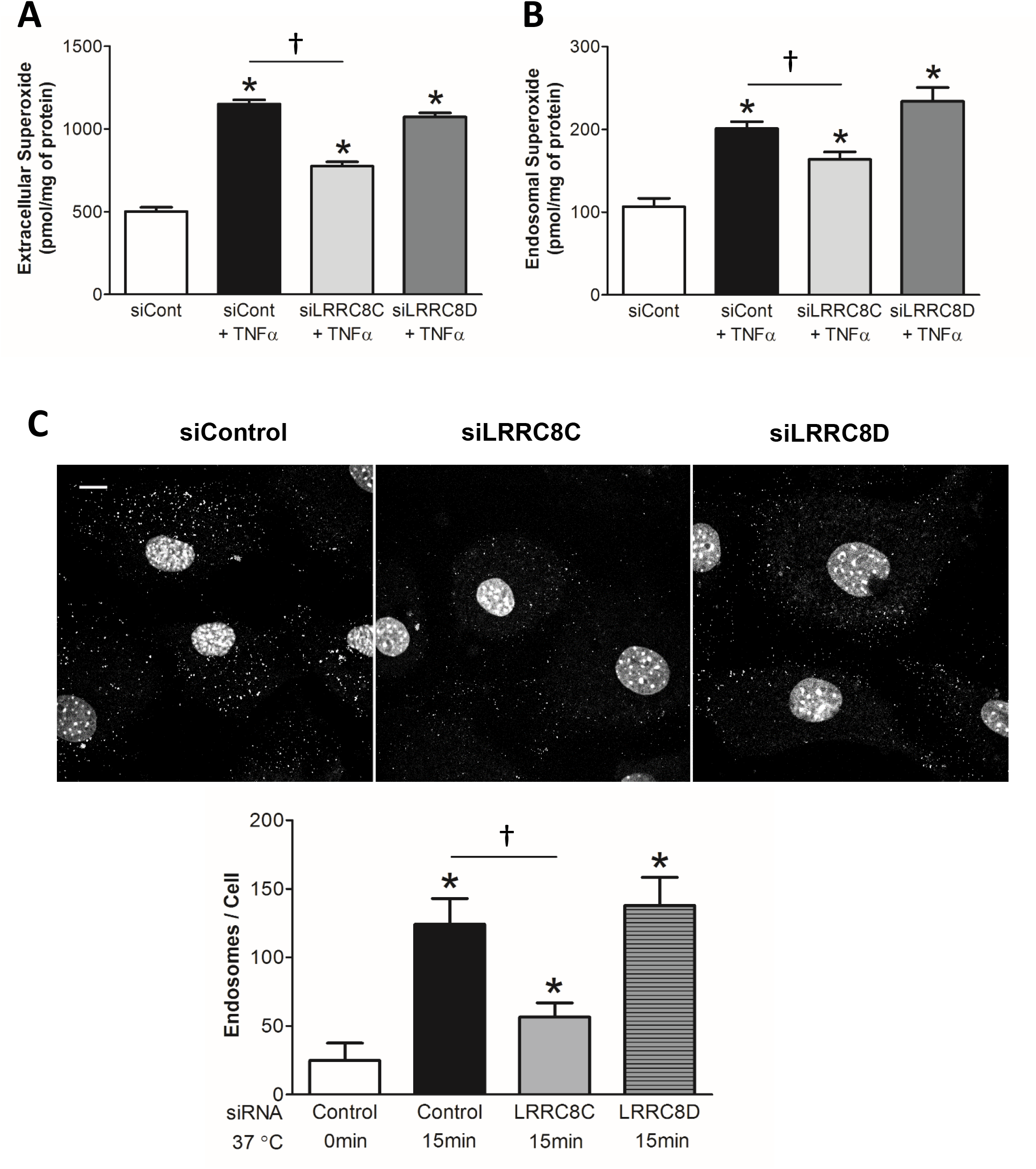
***A***, Extracellular superoxide production is increased in response to TNFα (CAT1H signal in media). This effect is inhibited by siLRRC8C but not by siLRRC8D. ***B***, Endosomal superoxide production is also increased in response to TNFα (CAT1H signal in cell fraction); this effect also was only inhibited by siLRRC8C. \**P* < 0.05 compared to siControl cells not exposed to TNFα, † *P* < 0.05 (n = 4). ***C***, Endocytosis of FITC-biotin-labelled TNFα is inhibited by siLRRC8C but not by siLRRC8D. Top: Typical images of fluorescent intracellular puncta in VSMCs treated for 15 mins with TNFα. Nuclei are stained with To-Pro-3. Scale (upper left): 10 µm. Bottom: Quantitation of puncta, expressed as endosomes/cell. * *P* < 0.05 compared to siControl cells not exposed to TNFα; † *P* < 0.05. Results are representative of at least three independent experiments.

TNFα receptor endocytosis is important for NF-κB activation (Choi *et al*., 2015). Impaired endosomal deposition of O_2_^•-^ could reflect reduced endosomal oxidase activity, as previously observed in cells lacking a ClC-3 transporter-mediated anion conductance (ClC-3) (Miller *et al*., 2007). However, a reduced CAT1H endosomal O_2_^•-^ signal could also result from impairment of the endocytic process. TNFα receptor endocytosis in VSMCs is O_2_^•-^-dependent, and was thus impaired in cells lacking Nox1 or LRRC8A (Choi *et al*., 2016). We assessed endocytosis of fluorescently labelled TNFα using biotin-labeled TNFα and FITC-avidin (Choi *et al*., 2015). The ability of murine VSMCs to form TNFα-containing endosomes was significantly reduced by siLRRC8C, while siLRRC8D had no effect (Fig. 3C).

### LRRC8 Expression in Human Tissues

Our siRNA data demonstrate that induced changes in LRRC8 isoform expression impact inflammatory cell signaling. However, the pathophysiological significance of these signaling changes depends on whether isoform expression is significant in tissues of interest and if it is altered by inflammatory disease states *in vivo*. Among 54 different tissues catalogued by the GTExPortal, LRRC8A expression is remarkably high in tissues containing an abundance of smooth muscle; the aorta (1^st^), tibial artery (2^nd^), coronary artery (6^th^), vagina (9^th^), and uterus (10^th^) rank in the top ten highest-expressing tissues (Fig. 4A). In the three blood vessels tested, the other LRRC8 isoforms, including 8C and 8D, are expressed at much lower copy number (<10% of LRRC8A), and their relative expression ranks much lower compared to LRRC8A (Fig. 4A).

**Figure 4.**
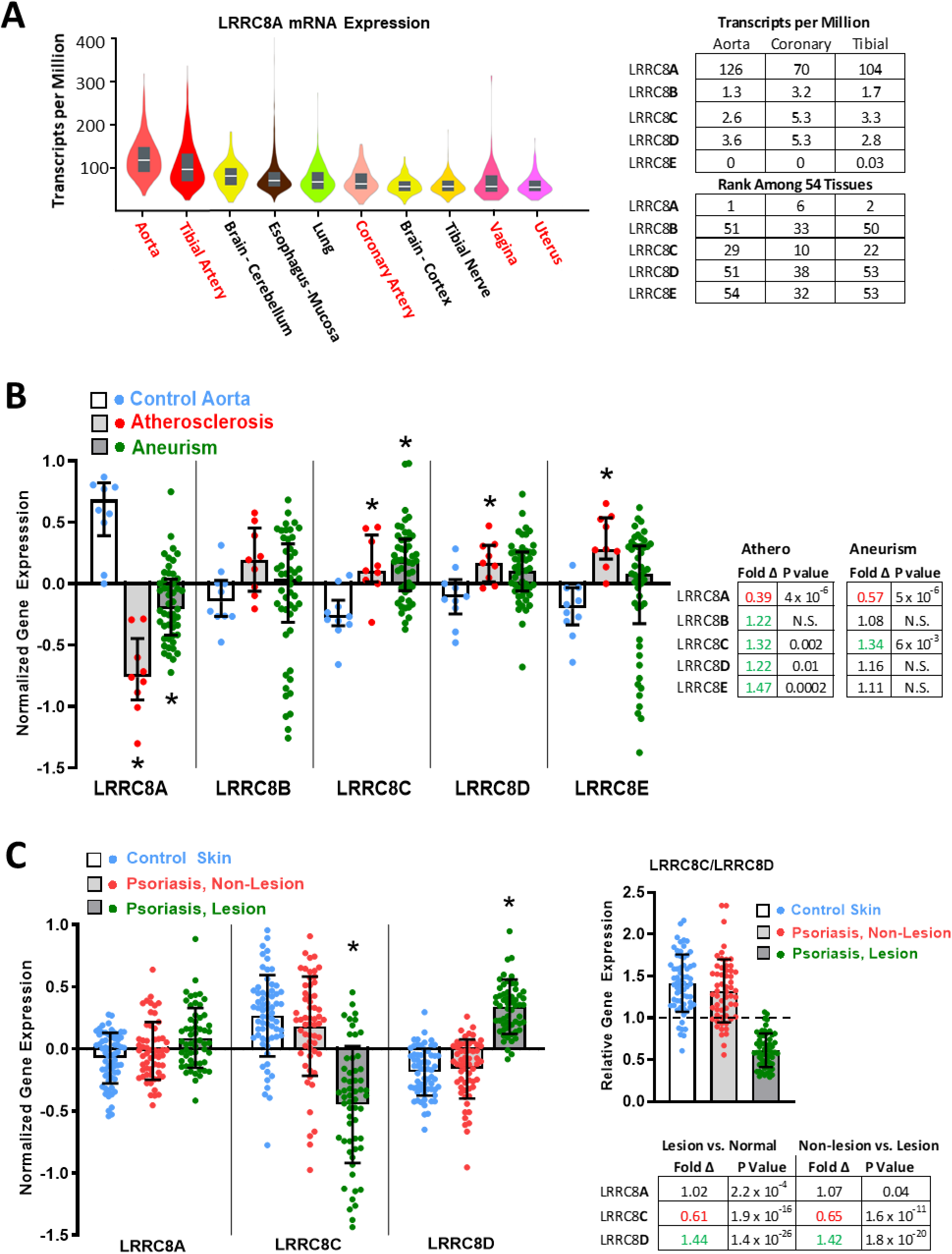
LRRC8 gene expression in health and disease. ***A***, Violin plots (left), including the median (white line) and interquartile range (grey bars), of LRRC8A expression (transcripts/million) in the 10 highest-expressing tissues of 54 human tissues assessed. LRRC8A is highly expressed in blood vessels and in other organ tissues with abundant smooth muscle (shown in red). The table to the right quantifies the relative expression of all LRRC8 isoforms (8A-8E) in the three blood vessels tested (top), and ranks the relative expression of each LRRC8 isoform in these vessels against expression in all 54 tissues (bottom). The n values vary between 142 for the uterus and 663 for the tibial artery. ***B***, Comparison of LRRC8 isoform expression in abdominal aortic biopsies from organ donors (Control, blue), and patients with either aortic atherosclerosis (red) or abdominal aneurisms (green). LRRC8A (ILMN_1739840), 8B (ILMN_1712128), 8C (ILMN2094396), 8D (ILMN_1763409) and 8E (ILMN_1669052) expression data are log_2_ transformed, such that a difference in normalized gene expression of 1.0 represents a doubling (or 50% decrease) in gene expression. LRRC8A expression is strongly reduced in both disease states. Fold differences and statistical significance of all changes are provided in the accompanying table (corrected P values obtained by GEO2R analysis). ***C***, Expression of LRRC8A (23347_at_s), 8C (228314_at) and LRRC8D (218684_at) was compared in skin biopsies from healthy patients (Control, n = 64), and from uninvolved (Non-lesion) and involved (Lesion) skin from psoriasis patients (n = 58). Psoriatic lesions demonstrate a small (7%) increase in LRRC8A expression, but remarkable changes in LRRC8C (decreased) and LRRC8D (increased). The shift in the ratio of LRRC8C to 8D expression in favor of LRRC8D was compared after 2^X^ transformation of the data (right). * P < 0.05.

We next compared LRRC8 isoform expression in biopsy samples from patients with inflammatory aortic disease or psoriasis. Both conditions are associated with significant TNFα-mediated inflammation (Chima & Lebwohl, 2018; Lamb *et al*., 2020). GSE57691 compares genome-wide expression in abdominal aortic biopsies from 10 healthy control patients (organ donors), 9 patients with aortic atherosclerosis, and 49 patients with abdominal aortic aneurisms (Biros *et al*., 2015). Fig. 4B displays transcript expression data for all LRRC8 isoforms. The most striking disease-related change is a reduction in LRRC8A expression in both atherosclerotic (61% decrease) and aneurismal vessels (43% decrease). By contrast, modest but statistically significant increases in expression occurred for LRRC8C, D and E in atherosclerosis, and for 8C in aneurismal disease. The relative expression of LRRC8C vs LRRC8D was not remarkably altered, as both increased to comparable degrees.

Psoriasis is an inflammatory skin disease that is largely driven by TNFα signaling, and is widely treated with selective anti-TNFα biologic agents (e.g. Etanercept) as first line therapy (Chima & Lebwohl, 2018). LRRC8A, 8C, and 8D gene expression was explored in GEO series GSE13355, which compares skin biopsies from healthy volunteers (control, n = 64) to both healthy (non-lesion) and inflamed (psoriatic lesion) skin from 58 patients with psoriasis (Nair *et al*., 2009). In contrast to aortic disease, LRRC8A expression is not meaningfully affected (increased 1.02-fold) in psoriatic lesions, but LRRC8C expression is significantly decreased, while 8D expression markedly increased (Fig 4C). When relative expression ratio (LRRC8C/LRRC8D) is compared under each condition, this ratio is positive in control and non-lesion groups (medians of 1.4 and 1.3) while in psoriatic lesions the ratio is reversed (median of 0.59).

### Chloramine-T inhibits native VRAC current and disrupts current block by DCPIB

We examined the oxidant sensitivity of VRAC currents in wild-type (WT) HEK293T cells and in LRRC8A^(-/-)^ mutant HEK293T cells expressing various LRRC8 constructs. All native LRRC8 isoforms were present in WT cells as assessed by RT-PCR, with 8D mRNA showing the most abundant expression among non-A isoforms (Fig. 5A, B). Expression levels of LRRC8B-E were not appreciably altered in 8A^(-/-)^ mutant cells nor was the abundance of LRRC8A message despite the presence of the inactivating mutation. In whole-cell recordings, application of hypotonic external saline (330 reduced to 255 mosm/L) to WT cells activated robust native VRAC currents that were nearly completely (>95%) blocked by 30 μM DCPIB (Fig. 5C, F). Comparable hypotonic current was absent in 8A^(-/-)^ mutant cells (Fig. 5D, G). Expression of LRRC8A-mCh in 8A^(-/-)^ mutant cells reconstituted DCPIB-sensitive hypotonic currents with largely WT behavior (Fig. 5E, G). Under isotonic conditions, WT cells typically (14/20 cells) displayed insignificant background currents (<6 pA/pF at +120 mV). Despite the use of pipette solution with 5% lower osmolality (315 mosm/L) than the control bath solution, a subset (6/20) of cells displayed significant initial currents (22 + 6 pA/pF; P < 0.001) that increased in amplitude with recording time (3 - 10 min) and resembled VRAC-mediated currents (Figure 6A). These isotonic currents were effectively blocked by DCPIB (Fig 6A, B), confirming that VRACs can be activated to varying degrees even under “isotonic” recording conditions. This finding is supported by our observations that inclusion of 10-20 μM external DCPIB reliably prevents activation of contaminating VRAC when recording other classes of anion current (J. Rohrbough, unpublished data). The addition of a C-terminal fluorescent tag to LRRC8 expression constructs constitutively activates current under isotonic conditions when all subunits express the tag (Gaitan-Penas *et al*., 2016). Accordingly, in 8A-mCh-expressing 8A^(-/-)^ cells, we recorded significantly larger isotonic currents (50 + 20 pA/pF at +120 mV, n = 3; Fig. 5E, G) compared to WT (10 + 3 pA/pF, n = 20; P < 0.02). These modest current densities nevertheless suggest that only minimal activation occurs for heteromeric channels containing both tagged and untagged subunits, and that hypotonic conditions are required for robust current activation.

**Figure 5.**
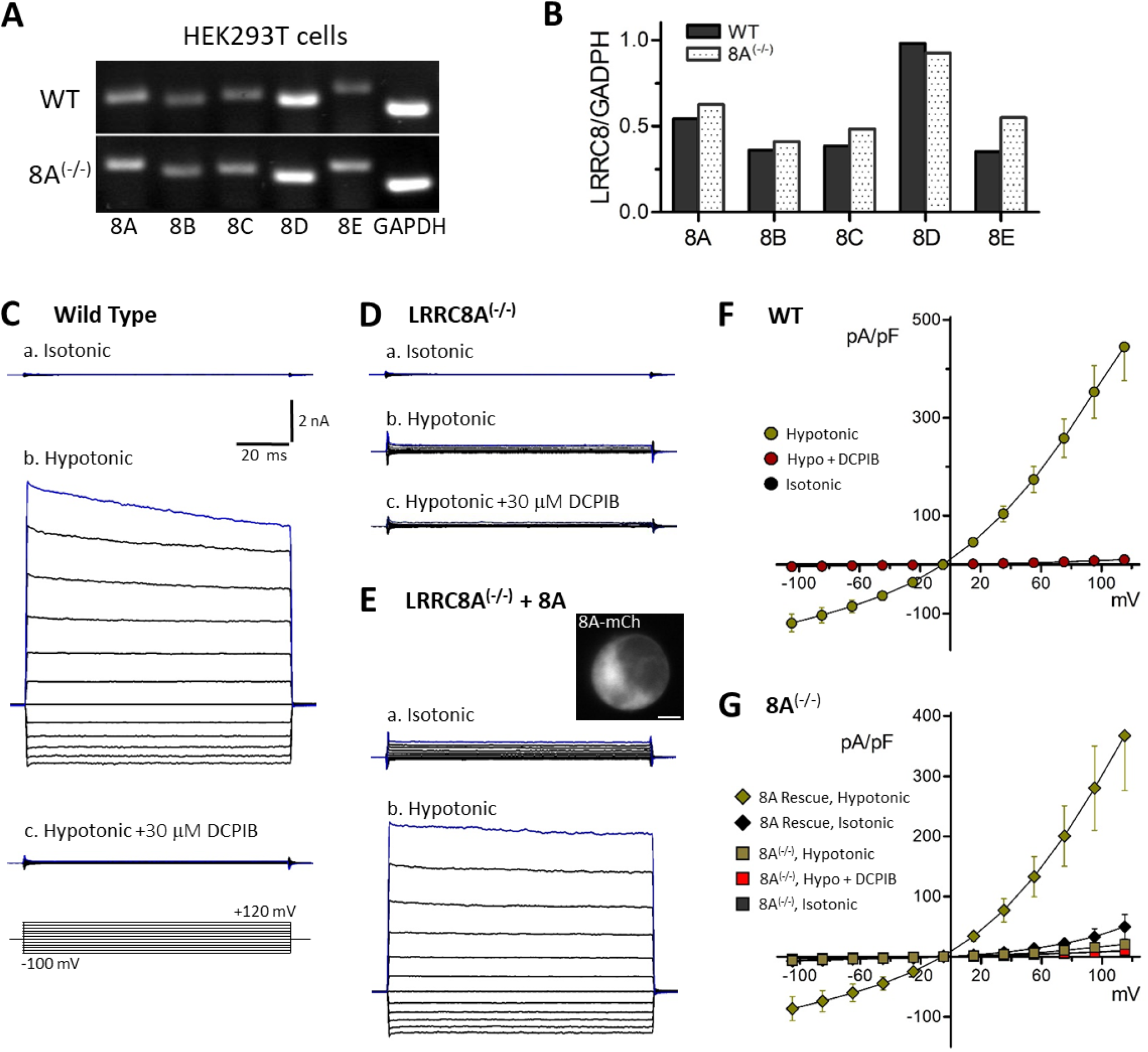
LRRC8A is required for endogenous VRAC current activation by hypotonic conditions. ***A***, Expression of LRRC8 isoforms (8A-8E) in WT (top) and LRRC8A^(-/-)^ HEK293 cells, as assessed by RT-PCR. For LRRC8A^(-/-)^, defective 8A mRNA is amplified by these primers but 8A protein is absent (data not shown). ***B***, Relative levels of LRRC8 isoform expression in HEK293 cells, revealing 8D to be the most abundantly expressed isoform. ***C-E***, Whole-cell currents recorded in HEK cells in response to 100 ms voltage test pulses (−100 to +120 mV). ***C***, Endogenous currents in a wild-type (WT) cell in isotonic (***a***, 330 msm/L) and hypotonic conditions (***b***, 255 mosm/L). DCPIB (***c***, 30 µM) potently blocks hypotonic current. The voltage pulse paradigm is depicted in ***c*** (bottom). Test pulses were preceded by a 10 ms prepulse to 0 mV from a holding potential of -40 mV. Scale: 2 nA, 20 ms (***C-E***). ***D***, In LRRC8A^(-/-)^ cells, hypotonic currents are virtually absent (***b***). ***E***, Expression of 8A-mCh in 8A^(-/-)^ cells results in small currents under isotonic conditions (***a***), and restores a robust response to hypotonic conditions (***b***). Inset image (***a***) shows 8A-mCh distribution in the recorded cell. ***F***, Current-voltage (I-V) relationships for wild-type cells. DCPIB blocks >95% of hypotonic-activated currents. ***G***, I-V relationships for LRRC8A^(-/-)^ mutant cells (square symbols) and 8A^(-/-)^ cells expressing 8A-mCh (Rescue; diamonds).

**Figure 6.**
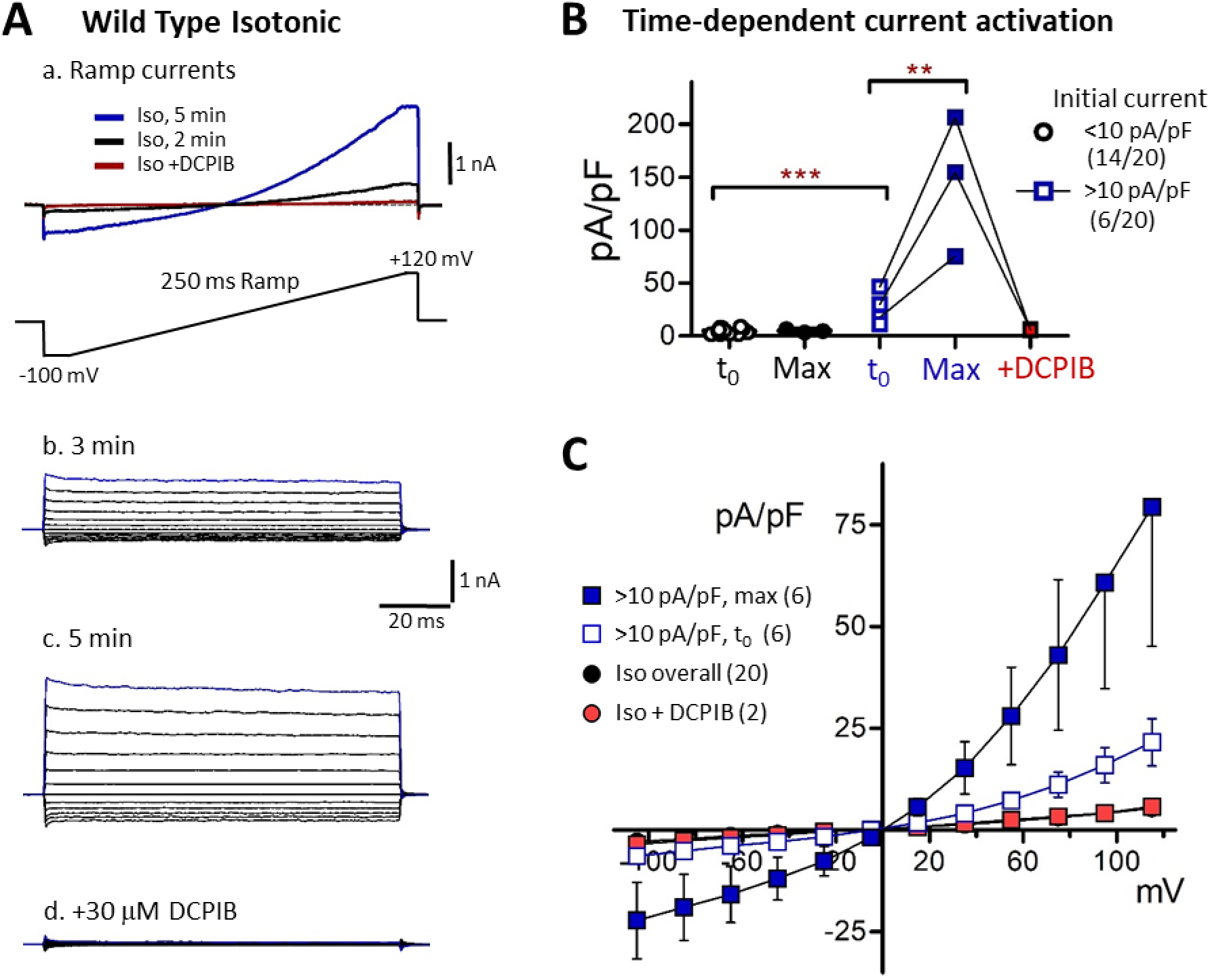
Time-dependent activation of endogenous VRAC currents in isotonic saline. ***A***, Time-dependent increase in VRAC currents in a wild-type HEK293 cells under isotonic conditions (330 mosm/L external, 315 mosm/L internal). ***a***, Currents induced by 250 ms voltage ramps (−100 to +120 mV; ramp protocol is depicted below) were increasingly activated over 5 mins after establishing whole cell configuration. Addition of 30 μM DCPIB completely blocked current. ***b-d***, Currents in response to voltage step-pulses (−100 to +120 mV), recorded at 3 min and 5 min following whole cell formation in the same cell. Qualitatively similar currents were observed under isotonic conditions in 6/20 WT recordings. ***B***, Quantified time-dependent changes in isotonic current amplitude (+120 mV) in two cells groups. The majority of cells (14/20; black symbols) exhibited small initial currents (<10 pA/pF) and showed no significant amplitude increase over 4-10 minutes (“max”: maximal time-dependent current). The remainder (6/20 cells; blue symbols) exhibited significantly larger initial currents (*** P < 0.0005 vs. majority group) with amplitudes that increased substantially over several minutes (** P < 0.005, max vs. t_0_), and were blocked by DCPIB (red symbols; connecting lines represent same-cell recordings). ***C***, Initial (t_0_; 1 - 2 min) and maximal (6 - 10 min) I-V relationships for VRAC-expressing cells (square symbols), compared to the overall I-V for all wild-type cells in isotonic conditions (circles).

When heteromerically expressed in *Xenopus* oocytes, LRRC8A/C and A/D channel currents were similarly inhibited by externally applied chloramine-T (ChlorT) (Gradogna *et al*., 2017). In HEK cells, externally applied ChlorT inhibited endogenous VRAC in WT cells (Fig. 7A), as well as reconstituted VRACs in 8A^(-/-)^ cells (Fig. 7A, B), with current amplitudes inhibited 50-60% by 1 mM ChlorT (Fig. 7C-E). Surprisingly, following ChlorT exposure, DCPIB was far less effective in blocking residual current (“post-ox. DCPIB”; Fig. 7Ab, Bb; 7E) compared to pre-oxidation conditions. This finding appears consistent with an oxidation-dependent change in the extracellular structure of VRACs as structural analysis indicated that DCPIB binds and blocks the outer channel pore of LRRC8A hexahomomers via a “cork-in-bottle” mechanism, interacting with Arg103 (Kern *et al*., 2019) located in extracellular loop 1 (EL1; Fig. 11A).

**Figure 7.**
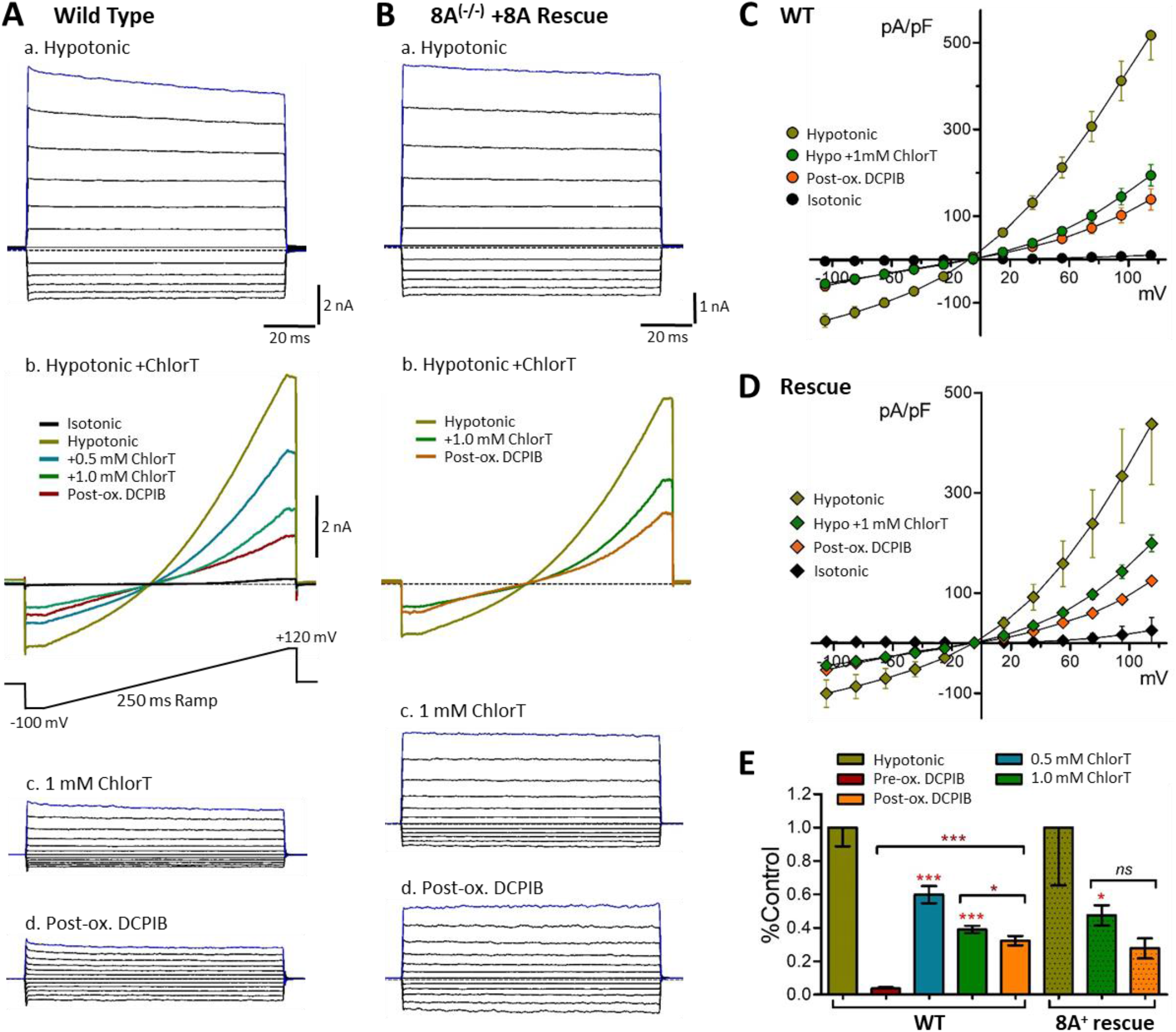
Endogenous VRAC currents are inhibited by the oxidant chloramine-T. ***A***, Wild-type (WT) HEK293 cells. ***a***, Currents in hypotonic conditions (−100 mV to +120 mV test pulses). Scale: 2 nA, 20 ms. ***b***, Responses in the same cell evoked by 250 ms voltage ramps (−100 mV to +120 mV; ramp protocol is depicted below) under the indicated sequential conditions (isotonic, hypotonic, hypotonic + ChlorT, hypotonic + 30 μM DCPIB). Current is progressively inhibited by 0.5 mM and 1 mM chloramine-T (ChlorT). Following ChlorT exposure, DCPIB block of residual current (post-ox DCPIB; red) is greatly reduced. Panels **c, d** show steady-state current families in 1 mM ChlorT and DCPIB. ***B***, LRRC8A knockout cell (LRRC8A^(-/-)^) expressing 8A-mCh (8A Rescue). ***a***, Currents evoked by voltage pulses under hypotonic conditions. ***b***, Ramp current responses in the same cell, in sequence (hypotonic, hypotonic + 1 mM ChlorT, hypotonic + 30 μM DCPIB). Panels **c, d** show steady-state currents in 1 mM ChlorT and DCPIB. ***C-D***, I-V relationships for wild-type (***C***) and 8A Rescue conditions (***D***), documenting the inhibition of hypotonic current by 1 mM ChlorT (n > 8 for wild type plots; n > 3 for rescue plots). ***E***, Summary of inhibitory effects of ChlorT and DCPIB on hypotonic ramp currents (+120 mV) for wild-type and 8A Rescue conditions. 1 mM ChlorT reduces wild-type peak current by 61% and 8A Rescue current by 53%. Post-oxidant DCPIB produces modest or insignificant block of residual current. Asterisks over bars denote significance level vs. hypotonic control amplitude (Paired ratio t-test; ***P < 0.0005; *P < 0.05). Brackets compare the indicated pair of bars (Paired ratio t-test or Mann-Whitney test). N > 7 for wild type; N = 3 - 4 for 8A rescue.

### LRRC8C and 8D currents display differential oxidant sensitivity

We hypothesized that oxidants produced by Nox1 can regulate activity of closely associated VRAC channels. Having established the association of Nox1 with LRRC8C and 8D, we examined ChlorT oxidant sensitivities of 8C vs. 8D channels, which has not previously been compared in mammalian cells. The trafficking of assembled VRAC channels to the plasma membrane is regulated by the LRRC8A intracellular loop (IL) sequence (P147-R262) (Yamada & Strange, 2018). 8C, D, and E chimeric proteins with the 8A IL sequence create functional homomeric channels in LRRC8 global knockout cells (Yamada & Strange, 2018). We expressed C-terminal tagged (monomeric Green (mGFP) or Cherry (mCh) fluorescent proteins) 8C-A IL and 8D-A IL chimeras in 8A^(-/-)^ cells. Under these conditions VRAC current is expected to also be mediated predominantly, if not exclusively, by homomeric 8C or 8D channels. In both 8C- and 8D-expressing cells, VRAC currents were substantially active under isotonic conditions, and further activated (∼2-fold) by hypotonic saline (Fig. 8A-B). LRRC8C currents were on average weakly inhibited by 1 mM ChlorT (23 + 4% reduction; n = 7; Fig. 8D). In contrast, 8D currents were potently inhibited by 0.5–1.0 mM ChlorT (Fig. 8Bc), displaying an over 85% reduction at 1 mM (Fig. 8E; P < 0.0005 vs. 8C, n = 6). We also assayed a tagged LRRC8A protein substituted with the EL1 sequence of LRRC8C that was also previously demonstrated to produce currents (Yamada & Strange, 2018). On average, currents in 8A-C EL1-expressing cells displayed no change in amplitude in response to 0.5 - 1.0 mM ChlorT (96 + 10% of control at 1 mM; n = 14; Fig. 8C, F). For each expression condition, addition of 30 μM DCPIB blocked currents present under either isotonic or hypotonic conditions by > 90% (Fig. 8D-F). This finding indicates that effective channel block by DCPIB does not strictly require the presence of Arg103 in LRRC8A since both 8C (lysine) and 8D (phenylalanine) have alternative residues at that position (Fig. 11B). However, as observed for endogenous currents, DCPIB was ineffective in blocking residual 8C or 8D currents following ChlorT exposure (Fig. 8G, H). In 8A-8C EL1-expressing cells, which were insensitive to ChlorT, post-ox DCPIB resulted in partial (∼40%) current block (Fig 8F, I). Oxidant-mediated structural changes to the extracellular channel domains therefore appears to disrupt DCPIB access or binding efficiency.

**Figure 8.**
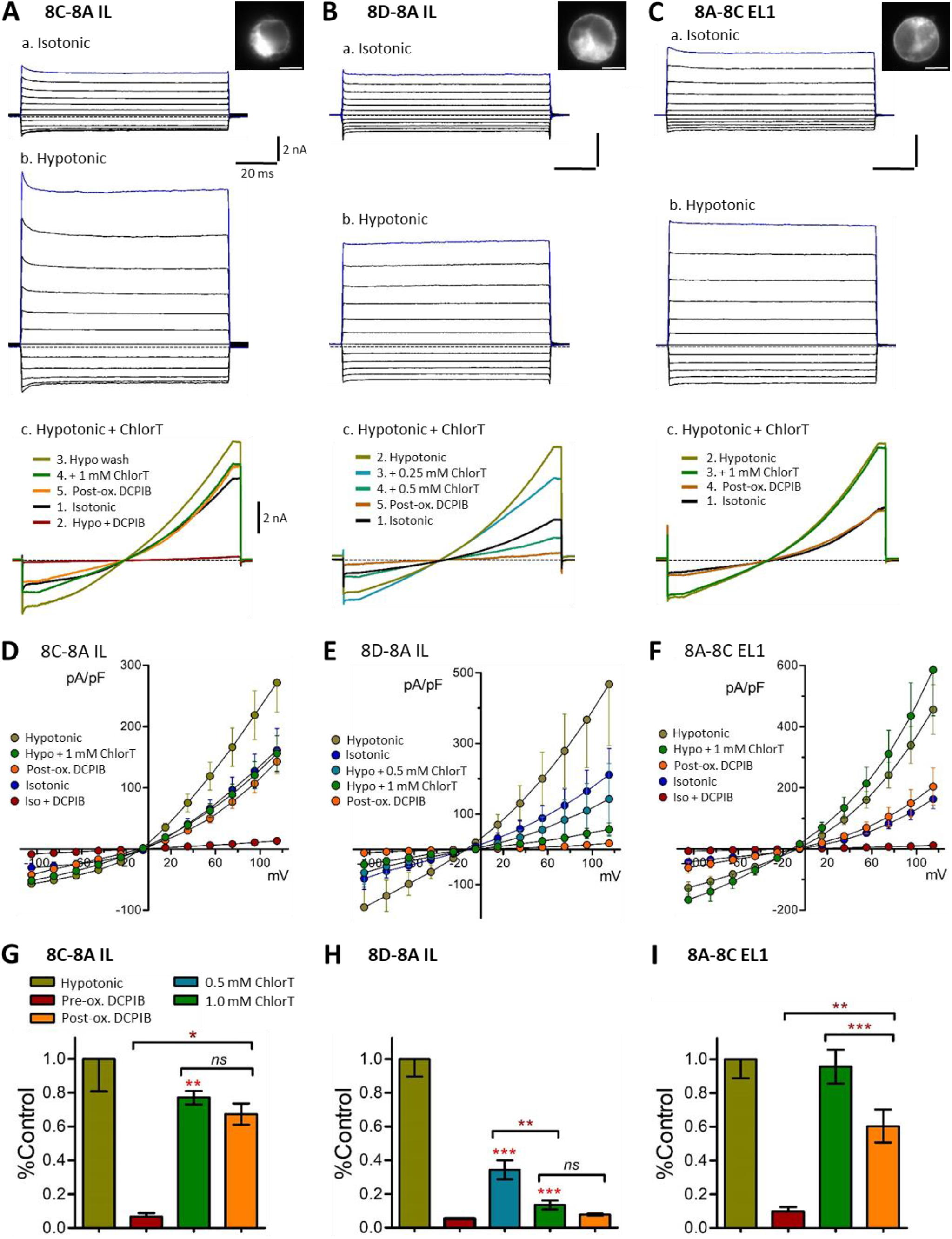
Differential oxidant sensitivity of homomeric LRRC8C, 8D and 8A channel currents. The 8A intracellular loop (IL) was substituted onto the 8C (***A***, 8C-8A-IL) and 8D (***B***, 8D-8A-IL) constructs tagged with mGFP or mCh and expressed in LRRC8A^(-/-)^ cells. Inset images (***a***) show construct fluorescence in the recorded cells. Panels ***a-b:*** Representative currents evoked by voltage pulses under isotonic (***a***) and hypotonic conditions (***b***). Scale: 2 nA, 20 ms (***A-C***). Panel ***c***: Current responses in individual cells to 250 ms voltage ramps (−100 to +120 mV) under the indicated series of conditions (isotonic, hypotonic, hypotonic + ChlorT, DCPIB). LRRC8C current is weakly inhibited by ChlorT (***Ac***), while 8D current (***Bc***) is strongly inhibited in a concentration-dependent manner by 0.25 and 0.5 mM ChlorT. ***C***, The first extracellular loop (EL1) of 8C was substituted onto LRRC8A-mGFP. Expressing cells display currents in both isotonic and hypotonic conditions (***a***,***b***). 8A-8C EL1 current is unaffected by 1 mM ChlorT (***Cc***). ***D-F***, Current-Voltage relationships for the three expression conditions, showing mean current densities evoked by 100 ms step pulses as in panels **a-b. *G-I***, Summary changes in ramp current amplitude (+120 mV) in response to ChlorT (0.5 - 1.0 mM) and DCPIB pre- and post-oxidant exposure. Asterisks over bars indicate significance vs. hypotonic control (Paired ratio t-test; *P < 0.05; **P < 0.005; ***P < 0.0005). Horizontal brackets compare the indicated pair of bars (paired ratio t-test or Mann-Whitney test)). 1 mM ChlorT weakly inhibits 8C (23% decrease, n = 7; ***G***), but potently inhibits 8D (>85% inhibition; P < 0.0005 vs. 8C, n = 6; ***H***). ChlorT has no net effect on 8A-8C EL1 current amplitude (n = 14; ***I***; see also Fig. 9E). Following oxidant exposure, DCPIB addition results in insignificant current block for 8C (n = 5) and 8D (n = 4), and only 40% block for 8A-8C EL1 (n = 10).

Endogenous VRAC currents are thought to be carried by heterohexamers composed of LRRC8A and one or more non-A subunits. We therefore also assessed oxidant responses in 8A^(-/-)^ cells co-expressing 8A-mCh with either 8C-mGFP (A/C) or 8D-mGFP (A/D) (Fig. 9). Both A/C and A/D coexpression conditions resulted in currents under isotonic conditions that were augmented by hypotonicity. 1 mM ChlorT moderately inhibited current in A/C-expressing cells (40 + 6% reduction; Fig. 9Ab, C, E), but exerted significantly stronger current inhibition in A/D-expressing cells (71 + 4% reduction; P < 0.005 vs A/C) (Fig. 9Bb, D, E). Currents in A/C and A/D-expressing cells likewise displayed insignificant post-oxidant DCPIB block (Fig. 9E). Based on the ability of LRRC8A-mCh expression to rescue current in 8A^(-/-)^ cells (Fig. 5), it is clear that the co-expressed 8A is able to combine with endogenous non-A protein(s) to contribute background VRAC current. However, the differential inhibition of currents in cells co-expressing A/C vs. A/D paralleled the behavior of the LRRC8C-A IL and 8D-A IL constructs.

**Figure 9.**
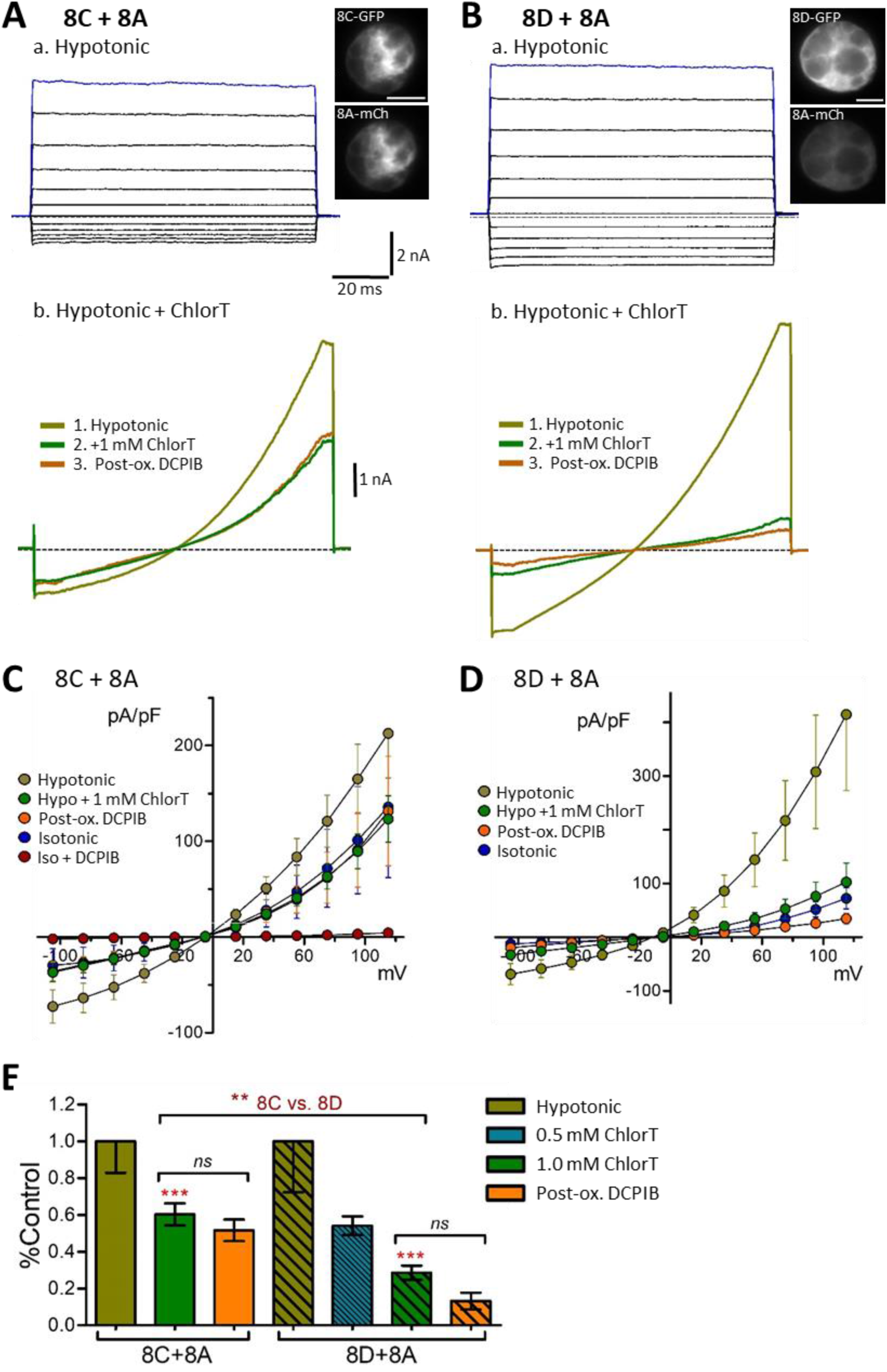
Differential oxidant sensitivity of 8C-8A and 8D-8A heteromeric channel currents. ***A, B***, Hypotonic current responses to voltage pulses (***a***) and 250 ms voltage ramps (***b***) in cells co-expressing 8C+8A (***A***) or 8D+8A (***B***). All isoforms were tagged with eGFP or mCh as described. Inset images (***panel***) show co-expressed fluorescence in the recorded cells. 1 mM ChlorT moderately inhibits current in 8C/A-expressing cells (***Ab***), and more strongly inhibits current in 8D/A-expressing cells (***Bb***). Post-oxidant DCPIB is ineffective in blocking residual currents. ***C, D***. I-V relationships for the indicated co-expression conditions. 8C+8A co-expression (***C***) resulted in substantial activation of DCPIB-sensitive currents in isotonic saline. ***E***, Summary of current response to ChlorT (0.5 - 1.0 mM) and pre/post DCPIB block. Asterisks over bars indicate significance vs. hypotonic control (Paired ratio t-test; **P < 0.005; ***P < 0.0005). Brackets compare the indicated pairs of bars (Paired Ratio t-test or Mann-Whitney test). N = 6 - 8 for 1 mM ChlorT; n = 4 - 6 for post-ox. DCPIB.

### Reducing agents exert minimal effects on ChlorT sensitive currents

The membrane-permeable reducing agent dithiothreitol (DTT) activated current in oocytes resulting from heteromeric LRRC8A/C or 8A/D channels, suggesting a baseline level of oxidation-dependent inhibition of these channels (Gradogna *et al*., 2017). However, DTT did not reverse current inhibition caused by ChlorT (Gradogna *et al*., 2017).

In WT HEK293 cells, application of DTT (10 mM, 4-5 min) had no consistent effect on endogenous current amplitudes under hypotonic conditions (111 + 30% of control amplitude, n = 5; Fig. 10A), and had no measurable effect on the efficacy of current block by 30 μM DCPIB (95 ± 2% block, n = 5; Fig. 10A). A membrane-impermeant reducing agent tris (2-carboxyethyl) phosphine (TCEP; 0.5 mM) likewise did not significantly alter WT hypotonic current amplitude (data not shown), and was ineffective in reversing current inhibition by 1 mM ChlorT (n = 2; Fig. 10B). However, TCEP application following ChlorT exposure appeared to partially restore sensitivity to DCPIB, which blocked ∼50% of the remaining current (P < 0.05, paired ratio t-test; Fig. 10B). We then examined isotonic current responses to TCEP in 8A^*(-/-)*^ cells expressing 8D-8A IL (Fig. 10C-E). TCEP exposure alone (0.5-1.0 mM, 2-5 min) did not significantly alter current amplitudes (92 ± 12% of control, n = 5; Fig. 10C), and subsequent DCPIB application blocked current by >90% (post-red. DCPIB, n = 3; Fig. 10C). ChlorT applied immediately following TCEP strongly inhibited current amplitudes (n = 2; Fig. 10C), indicating that ChlorT action was not impaired or slowed by a reduced environment. We conducted further tests to determine whether TCEP was able to reverse ChlorT-mediated current inhibition. ChlorT (1 mM, 1.5 - 3 min exposure) was first applied to induce current inhibition, followed by TCEP exposure (2.5 - 6 min) to assess reversibility (Fig. 10D). Current amplitudes demonstrated partial recovery in TCEP, ranging from 20%-30% to over 2-fold increase over the minimal ChlorT current level (66 ± 12% mean increase, n = 10; Fig. 10D, E). To test for current recovery attributable to ChlorT washout, we performed additional recordings that incorporated a wash period following ChlorT (Fig. 10D). We observed a comparable 50 ± 27% increase in current amplitudes associated with washout alone (n = 4; Fig. 10D, E); this increase did not differ statistically from the TCEP-associated increase (P > 0.7; Fig. 10E). TCEP addition following a wash period did not significantly increase current amplitude (Fig. 10D). We conclude that ChlorT-mediated changes underlying current inhibition are not readily reversible by a reducing agent under these experimental conditions.

**Figure 10.**
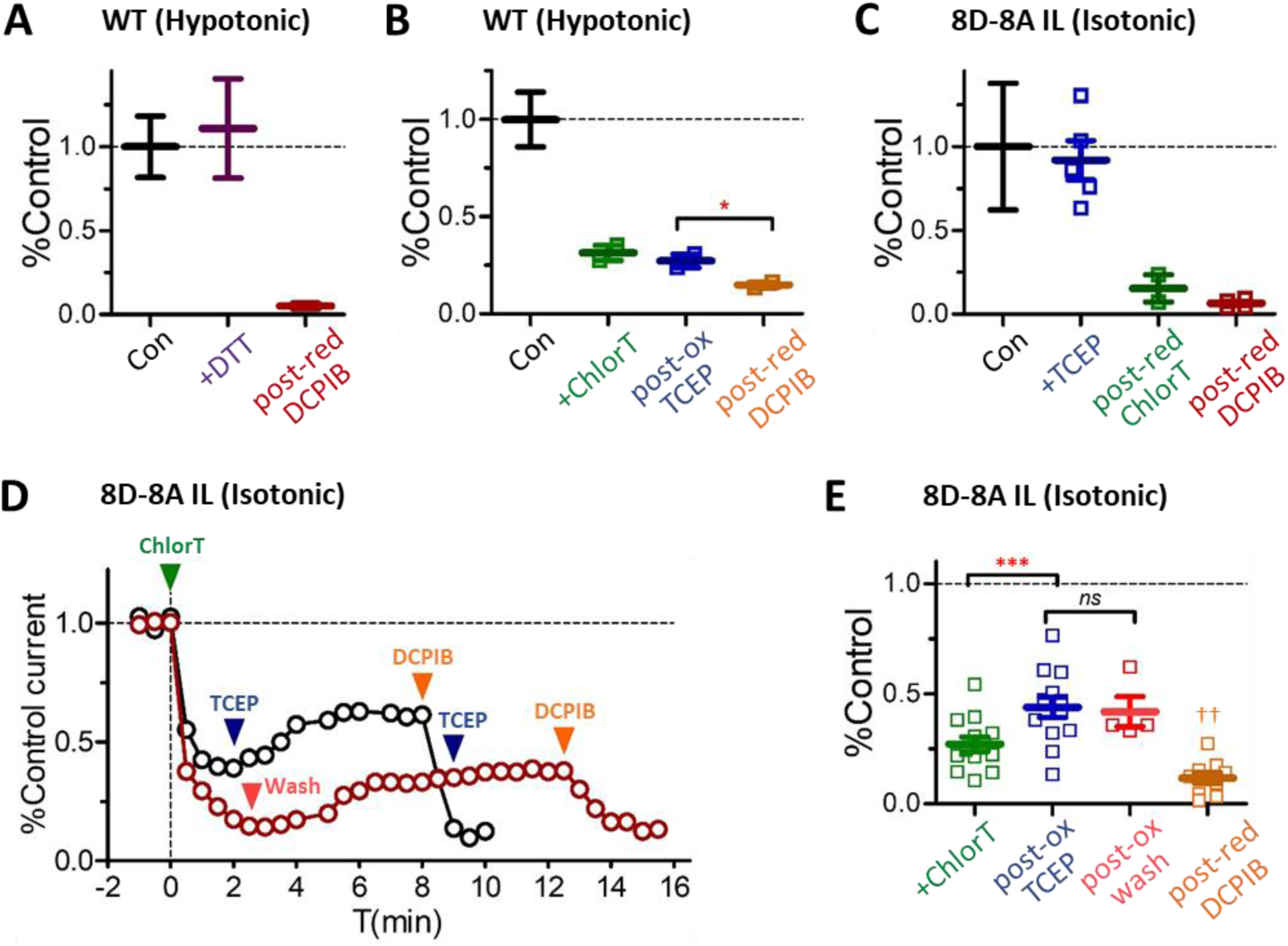
Wild-type and 8D channel currents are insensitive to externally applied reducing agents. ***A***, The membrane-permeant reductant dithiothretiol (DTT, 10 mM) does not significantly alter the amplitude of endogenous wild-type currents activated by hypertonic conditions (+120 mV; n = 5). Following DTT exposure, DCPIB (30 μM) addition results in nearly complete block of current (n = 3). ***B***, The membrane-impermeant reductant TCEP (0.5 mM) is ineffective in reversing ChlorT-mediated (1 mM) inhibition of wild-type currents. Subsequent addition of DCPIB results in significant additional inhibition (post-red DCPIB; * P < 0.05 vs. TCEP current level; Ratio paired t-test, n = 2). ***C-E***, Effects of TCEP application and ChlorT washout on isotonic currents in cells expressing 8D-8A IL. ***C***, TCEP (0.5-1.0 mM) exerted no effect on current amplitude (n = 5; symbols indicate individual recordings). Addition of 1 mM ChlorT following TCEP exposure produced strong inhibition (post-red ChlorT; n = 2), while DCPIB application following TCEP resulted in strong current block (post-red DCPIB; n = 3). ***D***, Current amplitude changes recorded in two cells (+120 mV). Exposure to 1 mM ChlorT at t_0_ (2-3 min exposure) resulted in rapid current inhibition by 60-80%. In cell 1 (black symbols), subsequent TCEP application (0.5 mM; 6 min) results in partial amplitude recovery, and addition of DCPIB results in >80% block of remaining current. Cell 2 (red symbols) displays a similar current amplitude recovery during ChlorT washout (red arrow, 6 min wash with control isotonic saline). Subsequent TCEP application (1 mM, 3 min) results in only minimal increase, while DCPIB blocks ∼70% of remaining current. ***E***, Summary of current changes during TCEP and wash experiments depicted in ***D***. Symbols indicate individual recordings; bold lines show average normalized current amplitudes. Green symbols: Current level following initial exposure to 1 mM ChlorT (n=13). Post-ox TCEP: 0.5 - 1.0 mM TCEP applied after ChlorT (no intervening wash; n=13). Current amplitude recovery following ChlorT washout (post-ox wash) was not distinguishable from the effects of TCEP (P > 0.7; n=4). Post-reductant DCPIB current level was assessed following sequential exposure to ChlorT and TCEP, with or without intervening wash (post-red DCPIB; n=9). Statistical comparisons were made using Paired ratio t-tests (*** P<0.0005; ^††^ P<0.01 vs. ChlorT).

**Figure 11.**
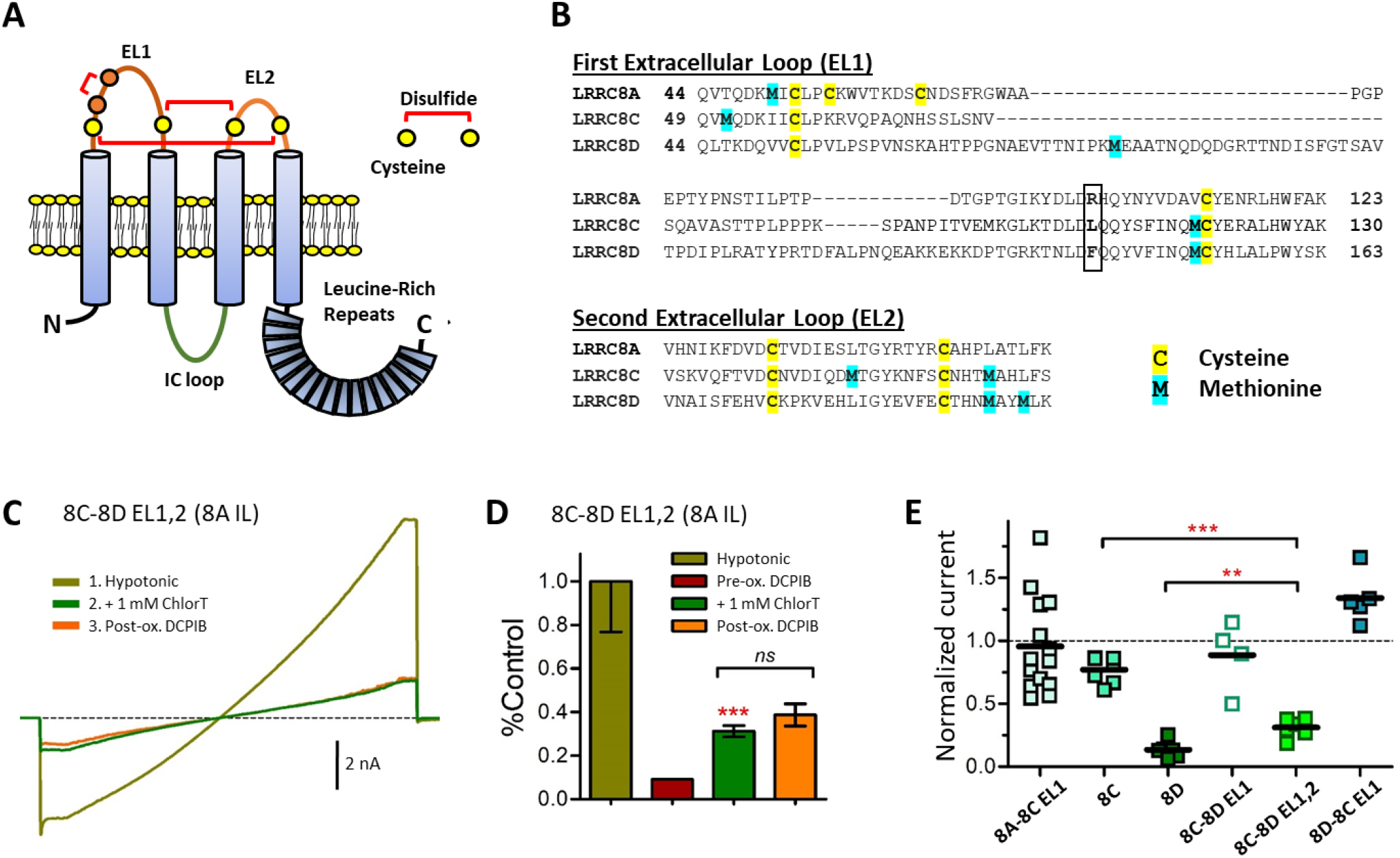
8D extracellular loop domains confer inhibitory ChlorT response to LRRC8C. ***A***, Structure of an LRRC8A monomer. The first extracellular loop (EL1) contains two conserved (C1, C4; yellow) and two non-conserved cysteines (C2, C3; orange). Conserved cysteines C1 and C4 form disulfide bonds with conserved cysteines in EL2, as shown. C2 and C3 form a third disulfide within EL1. The intracellular loop (IC; green) sequence (P147-R262) of LRRC8A was substituted for the homologous regions of LRRC8C and 8D for the experiments shown in Figs. 7 and 9. ***B***, EL1 and EL2 amino acid sequences for LRRC8A, C, and D, with potential redox-responsive cysteines (yellow) and methionines (blue) highlighted. R103 (boxed) of LRRC8A is implicated in DCPIB binding and block of LRRC8A homomers (Kern *et al*., 2019); 8C and 8D have lysine and phenyalanine, respectively, in the analogous position. ***C***, Currents evoked by 250 ms voltage ramps (−100 to +120 mV) in hypotonic saline from a cell expressing the chimeric LRR8C-8A IL protein with both 8D extracellular loops (8C-8D EL1,2). Current is strongly inhibited by 1 mM ChlorT. DCPIB is ineffective in blocking residual current following ChlorT exposure (post-ox. DCPIB). ***D***, Summary of 8C-8D EL1,2 current response to ChlorT and DCPIB. ChlorT inhibits current by 69% (n = 7; ***P < 0.0005 vs control hypotonic current; paired Ratio T-test). ***E***, Comparison of 1 mM ChlorT sensitivity among various 8A, 8C, and 8D chimeric constructs. Symbols represents normalized current amplitude relative to control hypotonic current in individual recordings, with mean values shown by horizontal lines. Substitution of 8C EL1 onto 8A (8A-8C EL1) confers no overall change in oxidant sensitivity. Homomeric 8C-8A IL shows weak inhibition, while 8D-8A IL shows strong inhibition. Substitution of 8D EL1 onto 8C (8C-8D EL1) does not alter the response to ChlorT. However, substitution of both EL1 and EL2 (8C-8D EL1,2) confers a strong inhibitory oxidant response (***P < 0.0006 vs. 8C; **P < 0.003 vs. 8D). Substitution of 8C EL1 onto 8D abolishes inhibitory responses.

### LRRC8D extracellular domains confer the inhibitory response to oxidation

To directly test the involvement of the extracellular structure of LRRC8D in ChlorT-mediated current inhibition, we created LRRC8D and 8C extracellular chimeras, targeting the EL1 and EL2 domains. The EL1 and EL2 amino acid sequences of LRRC8A, 8C and 8D and the general structural organization of LRRC8 proteins reveal multiple potential targets for redox active agents to mediate external structural changes to VRACs (Fig. 11A, B). In crystalized LRRC8A homohexamers, EL1 and EL2 are stabilized by three disulfide bonds (Deneka *et al*., 2018; Kasuya *et al*., 2018; Kern *et al*., 2019). In contrast to LRRC8A, the 8C and 8D EL domains each contain only 4 cysteines, yielding at most two disulfide bonds. The LRRC8C and 8D EL1 sequences differ significantly from each other and from 8A, with 8D EL1 containing 34 more residues compared to 8C. Methionine can also be oxidized by ChlorT, and methionine redox-dependent modulation can regulate protein function (Drazic & Winter, 2014). LRRC8C and 8D share a conserved methionine in each EL, while other methionine locations differ significantly between 8C, 8D, and 8A (Fig. 11B). Changes in the redox state of any of these sites could potentially regulate external channel structure, including the architecture of the DCPIB binding site in EL1 (Kern *et al*., 2019).

Substitution of the 8D EL1 sequence onto the LRRC8C-8A IL construct (8C-8D EL1) yielded currents that did not respond significantly to 1 mM ChlorT (11 ± 14% inhibition; P > 0.4 vs. 8C control; n = 4) (Fig. 11E). By contrast, substitution of both EL1 and EL2 of 8D onto 8C (8C-8D EL1,2; Fig. 11C, D) greatly enhanced current inhibition by ChlorT (69 ± 3% inhibition; n = 7; P < 0.0006 vs. 8C control; Fig. 11E). Conversely, substitution of 8C EL1 onto 8D abolished ChlorT-mediated inhibition and resulted in mild current augmentation (34 ± 9% increase; n = 5; P < 0.02 vs control) (Fig. 11E). Post-ox. DCPIB block was absent in each EL1 and EL1/EL2 variant (Fig. 11C-D, and data not shown). These results demonstrate that extracellular structural elements of both EL1 and EL1 of LRRC8D are required for the strong inhibition of currents by ChlorT. This raises the possibility that interaction between EL1 and EL2 is required for oxidant-dependent modulation, potentially disulfide formation between conserved cysteine residues or oxidation of multiple methionine residues.

## DISCUSSION

### Differential roles for LRRC8C and 8D in TNFα –dependent inflammatory signaling

Our results support the two-fold hypothesis that both LRRC8A/C and A/D channels physically interact with Nox1, but LRRC8A/C channels most efficiently support Nox1 oxidase activity. We show that LRRC8C knockdown depresses TNFα-dependent signaling, reducing TNFα-induced extracellular O_2_^•-^ production, TNFα receptor endocytosis, NF-κB activation and cell proliferation, as we previously established for LRRC8A. We therefore conclude that the TNFα inflammatory response is critically dependent on LRRC8A/C expression. In contrast, 8D knockdown does not significantly alter O_2_^-•^ production or receptor endocytosis, and enhances NF-κB activation. The pro-inflammatory role of the 8A/C channel may be related to its comparative insensitivity to oxidation, allowing persistent 8A/C current activity in an oxidized microenvironment that can support more sustained bursts of O_2_^•-^ production by Nox1. This conclusion is supported by the finding that LRRC8D channel currents are potently inhibited by oxidation, and this inhibitory sensitivity is conferred by the LRRC8D extracellular loop domains. In contrast to 8C, association of Nox1 with LRRC8A/D channels may facilitate more rapid and sensitive modulation of enzymatic activity via oxidative channel inhibition. In this model, LRRC8A/D channel oxidation acts as an “off switch”, providing negative feedback to ROS production and downregulating TNFα signaling (Fig. 12).

**Figure 12.**
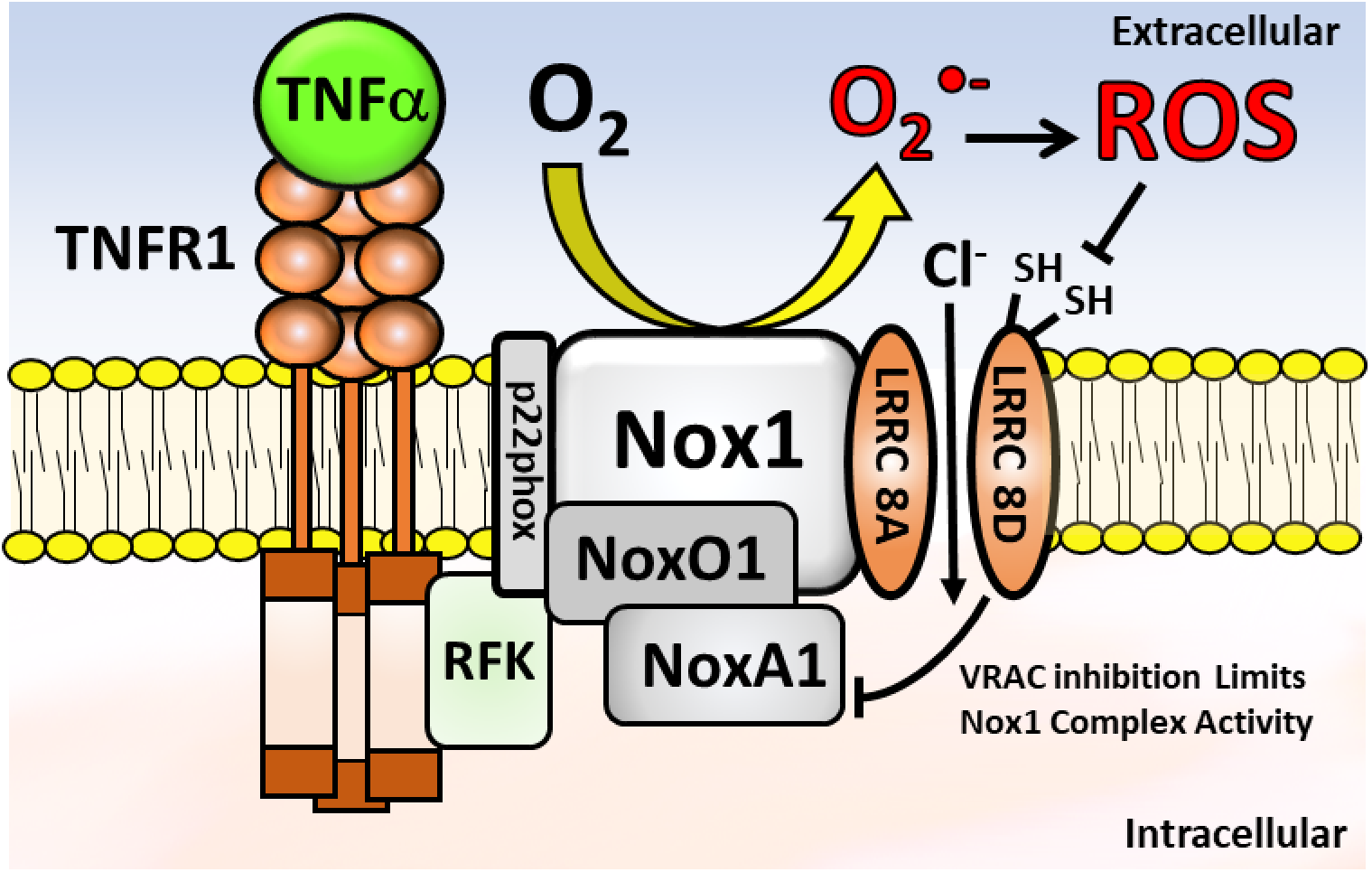
Working model of a multi-protein complex. The death domain of an occupied type 1 TNFα receptor (TNFR1, brown) associates with an assembled Nox1 complex (grey) via linkage to p22phox by riboflavin kinase (RFK, light green) (Yazdanpanah *et al*., 2009). The Nox1 complex also associates with LRRC8A (Choi *et al*., 2016), 8C and 8D. Activation of Nox1 by TNFα (bright green) leads to extracellular production of O_2_^•-^, which may itself act directly on LRRC8 VRACs. However, O_2_^•-^ also creates a variety of other reactive oxygen species (ROS) with the potential to oxidize redox-reactive sulfhydryl groups (−SH) on cysteine or methionine residues in the extracellular loops of LRRC8 VRACs. This results in inhibition of LRRC8A/D >> LRRC8A/C channels and thereby provides negative feedback regulation of Nox1.

VRACs are implicated in the control of proliferation, inflammation, and both apoptosis and autophagy-mediated cell death (Xia *et al*., 2016). DCPIB, which strongly inhibits VRAC current, is anti-inflammatory *in vivo*, reducing microglial inflammation and neuronal injury following cerebral ischemia (Han *et al*., 2014), and reducing infarct size after middle cerebral artery occlusion or hypoxia in rats and mice (Zhang *et al*., 2008; Alibrahim *et al*., 2013). We previously demonstrated that LRRC8A downregulation is an effective mechanism to reduce VRAC activity and negatively modulate Nox1-dependent TNFα signaling (Choi *et al*., 2016). Neuronal-specific LRRC8A knockout was protective in an ischemic stroke model (Zhou *et al*., 2020), and siLRRC8A has been shown to inhibit Angiotensin II-induced proliferation and migration in VSMCs (Lu *et al*., 2019). Both atherosclerosis (Kosmas *et al*., 2019) and aortic aneurismal disease (Rabkin, 2015) are thought to be significantly driven by a TNFα-dependent inflammatory process. We observed pronounced downregulation of LRRC8A expression in these diseases, but the LRRC8C/8D expression ratio remained stable. Decreased expression of a protein required for Nox1-dependent signaling is consistent with the Kyoto Encyclopedia of Genes and Genomes (KEGG) analysis of this gene expression dataset. Two of the three most downregulated pathways in atherosclerotic samples were cytokine-cytokine receptor interaction (hsa04060) which includes TNFα and IL-1β and receptors, and chemokine signaling (hsa04062) (Biros *et al*., 2015). Downregulation of these gene groupings, and of LRRC8A, is consistent with a compensatory response to a poorly controlled inflammatory state.

Relatively little is known regarding how LRRC8 isoforms traffic to the plasma membrane, or whether subunit association with LRRC8A is controlled by mechanisms other than relative subunit abundance. However, heteromeric A/E, A/D, and A/C channels have distinctly different biophysical properties (Konig & Stauber, 2019). Increased LRRC8D expression may result in decreased association of LRRC8A/C channels with Nox1, and *vice versa*, thereby impacting the response to Nox1-dependent cytokines. Psoriasis is a skin disease commonly treated by targeting TNFα signaling (Chima & Lebwohl, 2018). Inhibition of Nox1 reduced inflammation and improved human keratinocyte survival in an experimental model of psoriasis (Emmert *et al*., 2020). Thus, the observed expression changes favoring LRRC8D over LRRC8C in psoriatic skin are again consistent with downregulation of Nox-dependent inflammation. Changes in LRRC8 gene expression in inflamed tissues provide a potential link between these channels and human disease, and highlight the potential therapeutic utility of pharmacologic agents that selectively target VRACs.

Reactive oxygen production by NADPH oxidases is induced by cell swelling, and reactive oxygen species can be required for subsequent VRAC activation (Lambert, 2003; Holm *et al*., 2013). Signaling pathways such as epidermal growth factor and angiotensin II that activate Nox oxidases under isotonic conditions also activate VRACs (Browe & Baumgarten, 2004; Varela *et al*., 2004; Crutzen *et al*., 2012; Lu *et al*., 2019). This includes H_2_O_2_-dependent VRAC activation by TNFα in VSMCs (Matsuda *et al*., 2010). The LRRC8 subtype responsible for these currents is unknown. Currents mediated by heteromeric 8A/E channels in oocyte expression studies were activated by oxidants (Gradogna *et al*., 2017), supporting a hypothesis that native A/E channels may be activated by these signaling pathways. Similar results implicating A/E channel activation by oxidants have yet to be demonstrated in mammalian cells. However, ROS may also indirectly activate A/C or A/D currents via signals that are independent of any direct oxidant effect. In this scenario, the direct inhibitory effects of oxidants that we have described may act to buffer excessive VRAC activation. Importantly, Nox oxidases can deposit superoxide only into the extracellular space or within the lumen of intracellular vesicles (reviewed in (Lamb *et al*., 2020)). Although some ROS appear to be able to cross membranes, this requires specific pathways (ex. H_2_O_2_ moving through aquaporin channels (Bienert *et al*., 2007). Thus, it is critical to determine the specific site where oxidation acts on transmembrane proteins.

Heterologously expressed 8A/C and 8A/D heteromeric channels were previously reported to be inhibited by 1 mM ChlorT (Gradogna *et al*., 2017). We reasoned that the oxidized extracellular microenvironment created by Nox1 activation might exert a similar local inhibitory impact on closely associated VRAC channels in mammalian cells. LRR8D is clearly the most abundantly expressed native isoform in HEK293 cells, suggesting that native VRACs are composed primarily of the LRRC8A/D subtype. As in previous reports, we found that LRRC8A is an absolute requirement for VRAC current. Current could be fully reconstituted in LRRC8A null cells by heterologous expression of GFP-tagged LRRC8A, and both endogenous and reconstituted currents in HEK293 cells were similarly inhibited by 50%-60% by 1 mM ChlorT.

To compare isoform-specific responsiveness to oxidation, we utilized chimeric LRRC8C and D proteins substituted with the 8A-IL sequence, as well as LRRC8A-8C EL1, each previously shown to form functional homomeric channels (Yamada & Strange, 2018). In LRRC8A^(-/-)^ HEK cells, expression of these isoforms support currents under both isotonic and hypotonic conditions. We cannot exclude the possibility that these chimeric proteins also combine heteromerically with endogenous LRRC8 proteins. However, it is reasonable to assert that the recorded currents arise predominantly from “homomers” of the expressed chimeras. This conclusion is supported by the presence substantial current under isotonic conditions, which is consistent with the activating effect of the C-terminal fluorescent tag (Gaitan-Penas *et al*., 2016). LRRC8A, C, and D chimeric channel currents were further activated by hypotonicity, demonstrating that, as in oocytes, the C-terminal tag does not fully activate the channels. Notably, we showed that LRRC8D-8A IL currents are significantly more sensitive to inhibition by ChlorT than LRRC8C-8A IL currents, while LRRC8A-8C EL1 currents are unresponsive. We also demonstrated greater oxidant sensitivity for 8A/D vs. 8A/C currents in cells co-expressing LRRC8A-mCh with either 8D-GFP or 8C-GFP. Currents in co-expression experiments were unavoidably complicated by potential 8A-mCh combination with native non-A proteins. Despite this limitation, 8A/D coexpression resulted in significantly stronger (78%) current inhibition by ChlorTthan for 8A/C coexpression. These data support the results obtained with 8D and 8C chimera, and demonstrate that the enhanced oxidant sensitivity is not unique to homomeric LRRC8D-8A IL channels.

The crystal structure of LRRC8A homomers reveals two disulfide bonds between conserved cysteines which link EL1 with EL2, and a third between non-conserved cysteine residues within EL1 (Deneka *et al*., 2018; Kern *et al*., 2019). The status of these potential bonds (Fig. 11A, B) under *in vivo* conditions is completely unknown. In oocytes, expressed LRRC8A/C and A/D currents were similarly inhibited by ChlorT, but only LRRC8A/D currents were inhibited by MTSET, a membrane-impermeant modifier of extracellular cysteines (Gradogna *et al*., 2017). Substitution of both EL1 and EL2 of LRRC8D onto 8C was required to confer enhanced oxidant sensitivity (69% inhibition) strongly resembling 8D channel currents. This suggests that ChlorT must oxidize targets in both loops, or perhaps induce disulfide bond formation between the loops. ChlorT readily oxidizes glutathione (GSH) to a disulfide (GSSG), but cannot further oxidize cysteines to sulfinic or sulfonic acids (Shechter *et al*., 1975). ChlorT also can oxidize methionine residues, several of which are present in these loops (Fig. 11B), to potentially provide a mechanism of regulation (Drazic & Winter, 2014).

Membrane permeant (DTT) and impermeant (TCEP) reducing agents are theoretically capable of reversing these changes. However, these reagents did not alter LRRC8D VRAC currents, suggesting that in HEK293 cells either the extracellular structure of this channel is largely reduced, or that the redox active sites are not accessible to TCEP (Metcalfe *et al*., 2011). The *in vivo* extracellular redox potential (Cysteine/Cystine) is highly regulated, and becomes progressively more oxidized in cardiovascular disease and with aging, smoking, and obesity (Go & Jones, 2011). Thus, the “resting” redox state of LRRC8A/C and A/D channels *in vivo* may be variable. However, Nox1 activation is likely to effect a much more significant oxidation of the local environment than these baseline variations. We found that TCEP was unable to significantly reverse LRRC8D current inhibition following ChlorT exposure. This may indicate that key structural aspects of the oxidized extracellular loops are inaccessible, or cannot be “reset” by simple reduction. Disulfides can act as “switches” in the extracellular space, sensing redox potential and modulating protein function (Yi & Khosla, 2016). Oher redox-reactive extracellular proteins, such as thioredoxin or protein disulfide isomerases (PDI), can also potentially regulate these disulfides. PDI has previously been shown to associate with and modulate Nox1 activity intracellularly (Janiszewski *et al*., 2005), mediating Nox1 activation resulting by disulfide formation with p47phox, one of the cytoplasmic subunits of the oxidase (Gimenez *et al*., 2019).

A novel finding is that current block by DCPIB is impaired following ChlorT exposure. The potency of DCPIB as a VRAC blocker has been well established in numerous cell types. The effect of ChlorT on DCPIB block efficacy can be interpreted by considering the blockade mechanism. Cryo-EM imaging of LRRC8A homomers in lipid nanodisks has localized the DCPIB binding site to a ring formed by the association of R108 in EL1 of all six subunits (Kern *et al*., 2019). This ring is thought to form the ion selectivity filter of the channel (Deneka *et al*., 2018), though non-lysine residues in same positions are clearly capable of subserving the same function. Our results support the concept that the LRRC8 extracellular loop structure is modified by oxidation. The post-oxidant loss of DCPIB block efficacy occurred both for endogenous VRACs and in expressed channel constructs, although the effect was difficult to quantify for LRRC8D constructs (8D-8A IL and 8A/D co-expression) because ChlorT alone has such a profound inhibitory effect on these currents. Nevertheless, a reduction of post-oxidant DCPIB efficacy approached statistical significance in both cases. This supports the possibility of isoform-specific differences in oxidation-dependent modulation of DCPIB block. Post-oxidant DCPIB sensitivity was partially restored by TCEP exposure (Fig. 10D, E). In light of the potency of DCPIB block on native VRACs, these results support the idea that these channels are mostly in a reduced state under control conditions.

Inhibition of LRRC8A/D channels by extracellular oxidants may be relevant to many pathophysiologic conditions associated with oxidative stress (Fig. 12). In addition to the role of LRRC8 channels and ROS in vascular disease and psoriasis discussed above, LRRC8 VRACs have been broadly implicated in multiple types of cancer where they promote proliferation, migration and invasion. For this reason VRAC inhibitors have been proposed as novel targets for cancer therapy (reviewed in (Xu *et al*., 2020)). LRRC8A/D channels demonstrate significant permeability to uncharged osmolytes such as taurine, as well as to platinum-based anticancer drugs (Planells-Cases *et al*., 2015). An unbiased genome-wide screen identified both LRRC8A and D knockdown as inducers of resistance to platinum-based chemotherapeutic agents. Furthermore, low LRRC8D expression in ovarian cancer patients was correlated with a significant decrease in survival, suggesting that reduced uptake of these agents via LRRC8A/D channels reduced treatment effectiveness (Planells-Cases *et al*., 2015). Our results lend further relevance to these findings, given the near ubiquity of increased ROS production by cancer cells (Gorrini *et al*., 2013). LRRC8A/D channel oxidation in cancer cells may reduce uptake of chemotherapeutics. This problem may be amenable to targeted anti-oxidant therapy to enhance drug effectiveness (Ghoneum *et al*., 2020).

In summary, LRRC8A/C VRACs appear to be the primary LRRC8 isoform responsible for support of Nox1 activity and TNFα signaling. The relative abundance of the LRRC8C VRAC isoform has a significant influence on the inflammatory response of VSMCs to TNFα. These observations may have clinical relevance, as LRRC8 isoform expression is altered in people with atherosclerosis and psoriasis in a manner suggesting compensation for an inflamed state. The relative insensitivity of LRRC8A/C channel currents to inhibition by oxidation may render them better suited to support sustained Nox1 activity. These findings support the concept that LRRC8 VRACs are potential targets for anti-inflammatory therapeutic agents.

## ACKNOWLEDGMENTS

This project was supported by R01 HL128386 (F. Lamb) and by a Career Development Award from the American Heart Association 16SDG30610002 (H. Choi). We wish to thank Dr. Kevin Strange (Novo Biosciences and Vanderbilt University) for providing chimeric LRRC8 plasmids and Dr. Rajan Sah (Washington University) for the LRRC8A null HEK293 cells.

## ABBREVIATIONS

Ad: adenovirus
BCA: bicinchoninic acid
BSA: bovine serum albumin
CAT1H: 1-Hydroxy-2,2,6,6-tetramethylpiperidin-4-yl-trimethylammonium chloride
DCPIB: 4-(2-Butyl-6,7-dichloro-2-cyclopentyl-indan-1-on-5-yl) oxobutyric acid
DMEM: Dulbecco’s modified Eagle’s medium
eGFP: enhanced green fluorescent protein
ESR: electron spin resonance
FBS: fetal bovine serum
FITC: Fluorescein isothiocyanate
LRRC8: Leucine Rich Repeat Containing 8
NADPH: reduced nicotinamide-adenine dinucleotide phosphate
NF-κB: nuclear factor-kappa B
Nox: NADPH oxidase
ROS: reactive oxygen species
SEM: standard error of the mean
siRNA: small interfering ribonucleic acid
TNFα: tumor necrosis factor-α
TNFR: tumor necrosis factor-α receptor
VRAC: volume-regulated anion channel
VSMC: vascular smooth muscle cell

## REFERENCES

Alibrahim A, Zhao LY, Bae CY, Barszczyk A, Sun CL, Wang GL & Sun HS. (2013). Neuroprotective effects of volume-regulated anion channel blocker DCPIB on neonatal hypoxic-ischemic injury. Acta Pharmacol Sin 34, 113–118.

Bienert GP, Moller AL, Kristiansen KA, Schulz A, Moller IM, Schjoerring JK & Jahn TP. (2007). Specific aquaporins facilitate the diffusion of hydrogen peroxide across membranes. J Biol Chem 282, 1183–1192.

Biros E, Gabel G, Moran CS, Schreurs C, Lindeman JH, Walker PJ, Nataatmadja M, West M, Holdt LM, Hinterseher I, Pilarsky C & Golledge J. (2015). Differential gene expression in human abdominal aortic aneurysm and aortic occlusive disease. Oncotarget 6, 12984–12996.

Browe DM & Baumgarten CM. (2004). Angiotensin II (AT1) receptors and NADPH oxidase regulate Clcurrent elicited by beta1 integrin stretch in rabbit ventricular myocytes. J Gen Physiol 124, 273–287.

Chima M & Lebwohl M. (2018). TNF inhibitors for psoriasis. Semin Cutan Med Surg 37, 134–142.

Choi H, Dikalova A, Stark RJ & Lamb FS. (2015). c-Jun N-terminal kinase attenuates TNFalpha signaling by reducing Nox1-dependent endosomal ROS production in vascular smooth muscle cells. Free Radic Biol Med 86, 219–227.

Choi H, Ettinger N, Rohrbough J, Dikalova A, Nguyen HN & Lamb FS. (2016). LRRC8A channels support TNFalpha-induced superoxide production by Nox1 which is required for receptor endocytosis. Free Radic Biol Med 101, 413–423.

Choi H, Stark RJ, Raja BS, Dikalova A & Lamb FS. (2019). Apoptosis signal-regulating kinase 1 activation by Nox1-derived oxidants is required for TNFalpha receptor endocytosis. Am J Physiol Heart Circ Physiol 316, H1528–H1537.

Crutzen R, Shlyonsky V, Louchami K, Virreira M, Hupkens E, Boom A, Sener A, Malaisse WJ & Beauwens R. (2012). Does NAD(P)H oxidase-derived H2O2 participate in hypotonicity-induced insulin release by activating VRAC in beta-cells? Pflugers Arch 463, 377–390.

DeCoursey TE. (2016). The intimate and controversial relationship between voltage-gated proton channels and the phagocyte NADPH oxidase. Immunol Rev 273, 194–218.

DeCoursey TE, Morgan D & Cherny VV. (2003). The voltage dependence of NADPH oxidase reveals why phagocytes need proton channels. Nature 422, 531–534.

Deneka D, Sawicka M, Lam AKM, Paulino C & Dutzler R. (2018). Structure of a volume-regulated anion channel of the LRRC8 family. Nature.

Drazic A & Winter J. (2014). The physiological role of reversible methionine oxidation. Biochim Biophys Acta 1844, 1367–1382.

Emmert H, Fonfara M, Rodriguez E & Weidinger S. (2020). NADPH oxidase inhibition rescues keratinocytes from elevated oxidative stress in a 2D atopic dermatitis and psoriasis model. Exp Dermatol 29, 749–758.

Gaitan-Penas H, Gradogna A, Laparra-Cuervo L, Solsona C, Fernandez-Duenas V, Barrallo-Gimeno A, Ciruela F, Lakadamyali M, Pusch M & Estevez R. (2016). Investigation of LRRC8-Mediated Volume-Regulated Anion Currents in Xenopus Oocytes. Biophys J 111, 1429–1443.

Ghoneum A, Abdulfattah AY, Warren BO, Shu J & Said N. (2020). Redox Homeostasis and Metabolism in Cancer: A Complex Mechanism and Potential Targeted Therapeutics. International journal of molecular sciences 21.

Gimenez M, Schickling BM, Lopes LR & Miller FJ, Jr. (2016). Nox1 in cardiovascular diseases: regulation and pathophysiology. Clin Sci (Lond) 130, 151–165.

Gimenez M, Verissimo-Filho S, Wittig I, Schickling BM, Hahner F, Schurmann C, Netto LES, Rosa JC, Brandes RP, Sartoretto S, De Lucca Camargo L, Abdulkader F, Miller FJ, Jr. & Lopes LR. (2019). Redox Activation of Nox1 (NADPH Oxidase 1) Involves an Intermolecular Disulfide Bond Between Protein Disulfide Isomerase and p47(phox) in Vascular Smooth Muscle Cells. Arterioscler Thromb Vasc Biol 39, 224–236.

Go YM & Jones DP. (2011). Cysteine/cystine redox signaling in cardiovascular disease. Free Radic Biol Med 50, 495–509.

Gorrini C, Harris IS & Mak TW. (2013). Modulation of oxidative stress as an anticancer strategy. Nat Rev Drug Discov 12, 931–947.

Gradogna A, Gavazzo P, Boccaccio A & Pusch M. (2017). Subunit-dependent oxidative stress sensitivity of LRRC8 volume-regulated anion channels. J Physiol 595, 6719–6733.

Han Q, Liu S, Li Z, Hu F, Zhang Q, Zhou M, Chen J, Lei T & Zhang H. (2014). DCPIB, a potent volume-regulated anion channel antagonist, attenuates microglia-mediated inflammatory response and neuronal injury following focal cerebral ischemia. Brain Res 1542, 176–185.

Holm JB, Grygorczyk R & Lambert IH. (2013). Volume-sensitive release of organic osmolytes in the human lung epithelial cell line A549: role of the 5-lipoxygenase. Am J Physiol Cell Physiol 305, C48–60.

Janiszewski M, Lopes LR, Carmo AO, Pedro MA, Brandes RP, Santos CXC & Laurindo FRM. (2005). Regulation of NAD(P)H Oxidase by Associated Protein Disulfide Isomerase in Vascular Smooth Muscle Cells. J Biol Chem 280, 40813–40819.

Kasuya G, Nakane T, Yokoyama T, Jia Y, Inoue M, Watanabe K, Nakamura R, Nishizawa T, Kusakizako T, Tsutsumi A, Yanagisawa H, Dohmae N, Hattori M, Ichijo H, Yan Z, Kikkawa M, Shirouzu M, Ishitani R & Nureki O. (2018). Cryo-EM structures of the human volume-regulated anion channel LRRC8. Nat Struct Mol Biol 25, 797–804.

Kern DM, Oh S, Hite RK & Brohawn SG. (2019). Cryo-EM structures of the DCPIB-inhibited volume-regulated anion channel LRRC8A in lipid nanodiscs. Elife 8.

Konig B & Stauber T. (2019). Biophysics and Structure-Function Relationships of LRRC8-Formed Volume-Regulated Anion Channels. Biophys J 116, 1185–1193.

Kosmas CE, Silverio D, Sourlas A, Montan PD, Guzman E & Garcia MJ. (2019). Anti-inflammatory therapy for cardiovascular disease. Ann Transl Med 7, 147.

Lamb FS, Choi H, Miller MR & Stark RJ. (2020). TNFalpha and Reactive Oxygen Signaling in Vascular Smooth Muscle Cells in Hypertension and Atherosclerosis. American journal of hypertension.

Lamb FS, Moreland JG & Miller FJ, Jr. (2009). Electrophysiology of reactive oxygen production in signaling endosomes. Antioxid Redox Signal 11, 1335–1347.

Lambert IH. (2003). Reactive oxygen species regulate swelling-induced taurine efflux in NIH3T3 mouse fibroblasts. J Membr Biol 192, 19–32.

Liang W, Huang L, Zhao D, He JZ, Sharma P, Liu J, Gramolini AO, Ward ME, Cho HC & Backx PH. (2014). Swelling-activated Cl-currents and intracellular CLC-3 are involved in proliferation of human pulmonary artery smooth muscle cells. Journal of hypertension 32, 318–330.

Lu J, Xu F, Zhang J. (2019). Inhibition of angiotensin II-induced cerebrovascular smooth muscle cell proliferation by LRRC8A downregulation through suppressing PI3K/AKT activation. Human Cell 32, 316–325.

Matsuda JJ, Filali MS, Moreland JG, Miller FJ & Lamb FS. (2010). Activation of swelling-activated chloride current by tumor necrosis factor-alpha requires ClC-3-dependent endosomal reactive oxygen production. J Biol Chem 285, 22864–22873.

Metcalfe C, Cresswell P, Ciaccia L, Thomas B & Barclay AN. (2011). Labile disulfide bonds are common at the leucocyte cell surface. Open Biol 1, 110010.

Miller FJ, Jr., Filali M, Huss GJ, Stanic B, Chamseddine A, Barna TJ & Lamb FS. (2007). Cytokine activation of nuclear factor kappa B in vascular smooth muscle cells requires signaling endosomes containing Nox1 and ClC-3. Circ Res 101, 663–671.

Nair RP, Duffin KC, Helms C, Ding J, Stuart PE, Goldgar D, Gudjonsson JE, Li Y, Tejasvi T, Feng BJ, Ruether A, Schreiber S, Weichenthal M, Gladman D, Rahman P, Schrodi SJ, Prahalad S, Guthery SL, Fischer J, Liao W, Kwok PY, Menter A, Lathrop GM, Wise CA, Begovich AB, Voorhees JJ, Elder JT, Krueger GG, Bowcock AM, Abecasis GR & Collaborative Association Study of P. (2009). Genome-wide scan reveals association of psoriasis with IL-23 and NF-kappaB pathways. Nat Genet 41, 199–204.

Pedersen SF, Klausen TK & Nilius B. (2015). The identification of a volume-regulated anion channel: an amazing Odyssey. Acta Physiol (Oxf) 213, 868–881.

Planells-Cases R, Lutter D, Guyader C, Gerhards NM, Ullrich F, Elger DA, Kucukosmanoglu A, Xu G, Voss FK, Reincke SM, Stauber T, Blomen VA, Vis DJ, Wessels LF, Brummelkamp TR, Borst P, Rottenberg S & Jentsch TJ. (2015). Subunit composition of VRAC channels determines substrate specificity and cellular resistance to Pt-based anti-cancer drugs. EMBO J 34, 2993–3008.

Qian JS, Pang RP, Zhu KS, Liu DY, Li ZR, Deng CY & Wang SM. (2009). Static pressure promotes rat aortic smooth muscle cell proliferation via upregulation of volume-regulated chloride channel. Cell Physiol Biochem 24, 461–470.

Qiu Z, Dubin AE, Mathur J, Tu B, Reddy K, Miraglia LJ, Reinhardt J, Orth AP & Patapoutian A. (2014). SWELL1, a Plasma Membrane Protein, Is an Essential Component of Volume-Regulated Anion Channel. Cell 157, 447–458.

Rabkin SW. (2015). Accentuating and Opposing Factors Leading to Development of Thoracic Aortic Aneurysms Not Due to Genetic or Inherited Conditions. Front Cardiovasc Med 2, 21.

Shechter Y, Burstein Y & Patchornik A. (1975). Selective oxidation of methionine residues in proteins. Biochemistry 14, 4497–4503.

Syeda R, Qiu Z, Dubin AE, Murthy SE, Florendo MN, Mason DE, Mathur J, Cahalan SM, Peters EC, Montal M & Patapoutian A. (2016). LRRC8 Proteins Form Volume-Regulated Anion Channels that Sense Ionic Strength. Cell 164, 499–511.

Varela D, Simon F, Riveros A, Jorgensen F & Stutzin A. (2004). NAD(P)H oxidase-derived H(2)O(2) signals chloride channel activation in cell volume regulation and cell proliferation. J Biol Chem 279, 13301–13304.

Voss FK, Ullrich F, Munch J, Lazarow K, Lutter D, Mah N, Andrade-Navarro MA, von Kries JP, Stauber T & Jentsch TJ. (2014). Identification of LRRC8 heteromers as an essential component of the volume-regulated anion channel VRAC. Science 344, 634–638.

Xia Y, Liu Y, Xia T, Li X, Huo C, Jia X, Wang L, Xu R, Wang N, Zhang M, Li H & Wang X. (2016). Activation of volume-sensitive Cl-channel mediates autophagy-related cell death in myocardial ischaemia/reperfusion injury. Oncotarget 7, 39345–39362.

Xu R, Wang X & Shi C. (2020). Volume-regulated anion channel as a novel cancer therapeutic target. Int J Biol Macromol 159, 570–576.

Yamada T & Strange K. (2018). Intracellular and extracellular loops of LRRC8 are essential for volume-regulated anion channel function. J Gen Physiol.

Yazdanpanah B, Wiegmann K, Tchikov V, Krut O, Pongratz C, Schramm M, Kleinridders A, Wunderlich T, Kashkar H, Utermohlen O, Bruning JC, Schutze S & Kronke M. (2009). Riboflavin kinase couples TNF receptor 1 to NADPH oxidase. Nature 460, 1159–1163.

Yi MC & Khosla C. (2016). Thiol-Disulfide Exchange Reactions in the Mammalian Extracellular Environment. Annu Rev Chem Biomol Eng 7, 197–222.

Zhang Y, Zhang H, Feustel PJ & Kimelberg HK. (2008). DCPIB, a specific inhibitor of volume regulated anion channels (VRACs), reduces infarct size in MCAo and the release of glutamate in the ischemic cortical penumbra. Exp Neurol 210, 514–520.

Zhou JJ, Luo Y, Chen SR, Shao JY, Sah R & Pan HL. (2020). LRRC8A-dependent volume-regulated anion channels contribute to ischemia-induced brain injury and glutamatergic input to hippocampal neurons. Exp Neurol 332, 113391.

